# Role of a holo-insertase complex in the biogenesis of biophysically diverse ER membrane proteins

**DOI:** 10.1101/2023.11.28.569054

**Authors:** Katharine R. Page, Vy N. Nguyen, Tino Pleiner, Giovani Pinton Tomaleri, Maxine L. Wang, Alina Guna, Ting-Yu Wang, Tsui-Fen Chou, Rebecca M. Voorhees

## Abstract

Mammalian membrane proteins perform essential physiologic functions that rely on their accurate insertion and folding at the endoplasmic reticulum (ER). Using forward and arrayed genetic screens, we systematically studied the biogenesis of a panel of membrane proteins, including several G-protein coupled receptors (GPCRs). We observed a central role for the insertase, the ER membrane protein complex (EMC), and developed a dual-guide approach to identify genetic modifiers of the EMC. We found that the back of sec61 (BOS) complex, a component of the ‘multipass translocon’, was a physical and genetic interactor of the EMC. Functional and structural analysis of the EMC•BOS holocomplex showed that characteristics of a GPCR’s soluble domain determine its biogenesis pathway. In contrast to prevailing models, no single insertase handles all substrates. We instead propose a unifying model for coordination between the EMC, multipass translocon, and Sec61 for biogenesis of diverse membrane proteins in human cells.

## INTRODUCTION

Integral membrane proteins are essential across all biological systems, including in mammalian cells and their pathogens. Human membrane proteins mediate a range of processes from cell-to-cell signaling to metabolite transport (von Heijne, 2007). Similarly, many viruses encode membrane proteins that are critical for fusion with a host cell, organization of the replication machinery, and transport of ions and small molecules (by viroporins) that enhance infectivity and morbidity (Harrison, 2008; Lenard, 2008). In order to carry-out these functions, both the transmembrane (TMDs) and soluble domains have evolved diverse charge, hydrophobicity, and length (von Heijne, 2007). The accurate insertion and folding of these topologically and biophysically diverse proteins therefore represents a major challenge in human cells. Despite the importance of this process, how cells regulate biogenesis of the full complexity of the mammalian and viral membrane proteome is not understood.

The majority of membrane proteins, destined for either the plasma membrane or secretory system, begin their biogenesis at the endoplasmic reticulum (ER) (Rapoport et al., 1996; Shao & Hegde, 2011). For multipass proteins, a nascent polypeptide is captured in the cytosol by the signal recognition particle (SRP) for delivery of the ribosome nascent chain complex to the membrane (Halic & Beckmann, 2005; Shan & Walter, 2005). Once at the ER, TMDs must be inserted into the lipid bilayer. Insertion requires two simultaneous processes: (i) transfer of the hydrophobic TMD from the aqueous cytosol to a membrane-spanning topology within the lipid bilayer, and (ii) translocation of an associated soluble domain across the membrane into the ER lumen. The latter of which is energetically unfavorable and therefore typically catalyzed by a membrane protein insertase in cells (Guna, Hazu, et al., 2023).

The textbook model posits that the Sec61 translocation channel is the major insertase for multipass membrane proteins. It was hypothesized that its unique clam-shell architecture could accommodate all aspects of membrane protein biogenesis: axial opening creates a pore in the membrane for translocation into the ER lumen, while lateral opening would permit partitioning of a TMD into the bilayer (Van den Berg et al., 2004; Voorhees & Hegde, 2016). Studies of signal sequences suggested that opening of Sec61 is triggered by binding of a sufficiently hydrophobic segment of the nascent chain at the lateral gate of the channel (Rapoport et al., 2017). However, many multipass membrane proteins contain poorly hydrophobic TMDs, which while required for function, cannot autonomously gate Sec61 (Enquist et al., 2009; Schorr et al., 2020). Therefore, a simple model in which each TMD is sequentially inserted into the lipid bilayer by Sec61 alone cannot explain the insertion or folding of most multipass membrane proteins.

Recently, it has instead been proposed that substrates, including the physiologically essential family of G-protein coupled receptors (GPCRs), are inserted by a ‘multipass translocon’ that uses Sec61 as a structural scaffold, but does not rely on its insertase activity (McGilvray et al., 2020; Smalinskaitė et al., 2022; Sundaram et al., 2022). The multipass translocon is a dynamic, hetero-oligomeric 8-subunit complex that includes the GET and EMC like (GEL), back of Sec61 (BOS), and PAT complexes (Figure S1A). The GEL complex— composed of the Oxa1 superfamily insertase, TMCO1 and its binding partner, OPTI—serves as the dedicated insertase of the multipass translocon, and is postulated to integrate nascent TMDs as they emerge from the ribosome behind Sec61. The PAT complex, containing Asterix and CCDC47, has two proposed roles: Asterix chaperones hydrophilic TMDs within the lipid bilayer during synthesis of a multipass protein (Chitwood & Hegde, 2020; Meacock et al., 2002); while CCDC47 directly engages the ribosome and closes the lateral gate of Sec61, helping to guide TMDs to the multipass translocon (Smalinskaitė et al., 2022; Sundaram et al., 2022). Finally, the function of the BOS complex remains unknown, but is thought to act as scaffold for recruitment of the remaining multipass components (Smalinskaitė et al., 2022; Sundaram et al., 2022). Together, these factors create a protected lipid cavity behind Sec61 thought to facilitate multipass membrane protein insertion and folding. In support of this model, it was shown that biogenesis of the GPCR rhodopsin is unaffected by inhibitors that prevent access to the lateral gate of Sec61, and nascent TMDs crosslink to multipass translocon components as they emerge from the ribosome. However, whether the GEL complex, which is metazoan specific and not essential in humans (Karczewski et al., 2020), is responsible for the integration of all multipass TMDs remains unclear.

Indeed, earlier work established that an additional insertase, the ER membrane protein complex (EMC), was required for biogenesis of many multipass membrane proteins. In mammals, the EMC is an abundant, nine-subunit complex that functions as both an insertase and chaperone (Chen et al., 2023; Guna et al., 2018; Shurtleff et al., 2018; Tian et al., 2019). In addition to post-translational insertion of a subset of tail anchored proteins, the EMC also co-translationally inserts the first TMD of many GPCRs and other multipass membrane proteins that position their N-terminus in the ER lumen or extracellular environment (i.e. adopt an N_exo_ topology) (Chitwood et al., 2018). Indeed, expression of rhodopsin, which does not rely on the lateral gate of Sec61 for insertion, is EMC dependent (Satoh et al., 2015). However, the function of all nine of EMC’s subunits, in particular those that form its large lumenal domain that is not directly involved in insertion, is not known.

Structures of the yeast and human EMC show that substrate TMDs are inserted into the bilayer via a positively charged hydrophilic groove, through which the substrate’s soluble N-terminus must also translocate (Bai et al., 2020; Miller-Vedam et al., 2020; O’Donnell et al., 2020; Pleiner et al., 2020). The positioning of positively charged residues within the membrane is a conserved feature of the Oxa1 superfamily of insertases and is required for their activity (Borowska et al., 2015; Kumazaki et al., 2014; McDowell et al., 2020; Pleiner et al., 2020, 2023). In contrast to the prevailing model, it is likely that multipass substrates are therefore directly delivered by SRP to the EMC (Wu & Hegde, 2023), leaving the EMC to act upstream of Sec61 and the multipass translocon.

However, this model leaves several central unanswered questions for how human and viral membrane proteins are accommodated by the biogenesis and quality control machinery in the ER. First, whether or how the EMC coordinates with the multipass translocon during multipass biogenesis is not known. Second, if the EMC is responsible for insertion of the first N_exo_ TMD of many membrane proteins (including GPCRS), how substrates are transferred between the EMC, Sec61, and the multipass translocon is not clear. Finally, a systematic analysis of the substrate specificity and cooperation of the suite of biogenesis factors in the ER to ensure insertion and folding of their diverse clients, has not been explored.

## RESULTS

### Systematic analysis of membrane protein biogenesis

With the goal of unbiasedly identifying factors required for biogenesis of diverse membrane proteins, we selected a panel of substrates with distinct topologies, biophysical properties, and number of TMDs (Figure 1 and S1B). In this initial panel we included the co-translationally targeted multipass proteins AGTR2 and the viral ORF3a and M from SARS-CoV-2, as well as the post-translationally targeted single spanning tail-anchored protein, Sec61β. AGTR2 is a seven-TMD G-protein coupled receptor (GPCR; Figure 1A). GPCRs represent the largest single family of membrane proteins encoded by the human genome and are responsible for physiologically important signaling processes throughout the human body (Heldin et al., 2016; O’Hayre et al., 2014). AGTR2, like most GPCRs, contains TMDs with varying hydrophobicities, including those predicted to insert autonomously and those likely to only insert in the context of the intact multispanning protein (Cvicek et al., 2016). Outside of the TMDs, AGTR2 contains small cytosolic and extracellular loops, and a neutrally charged N-terminal soluble domain that must be translocated across the ER membrane (i.e. adopts an N_exo_ topology) during biogenesis.

**Figure 1.**
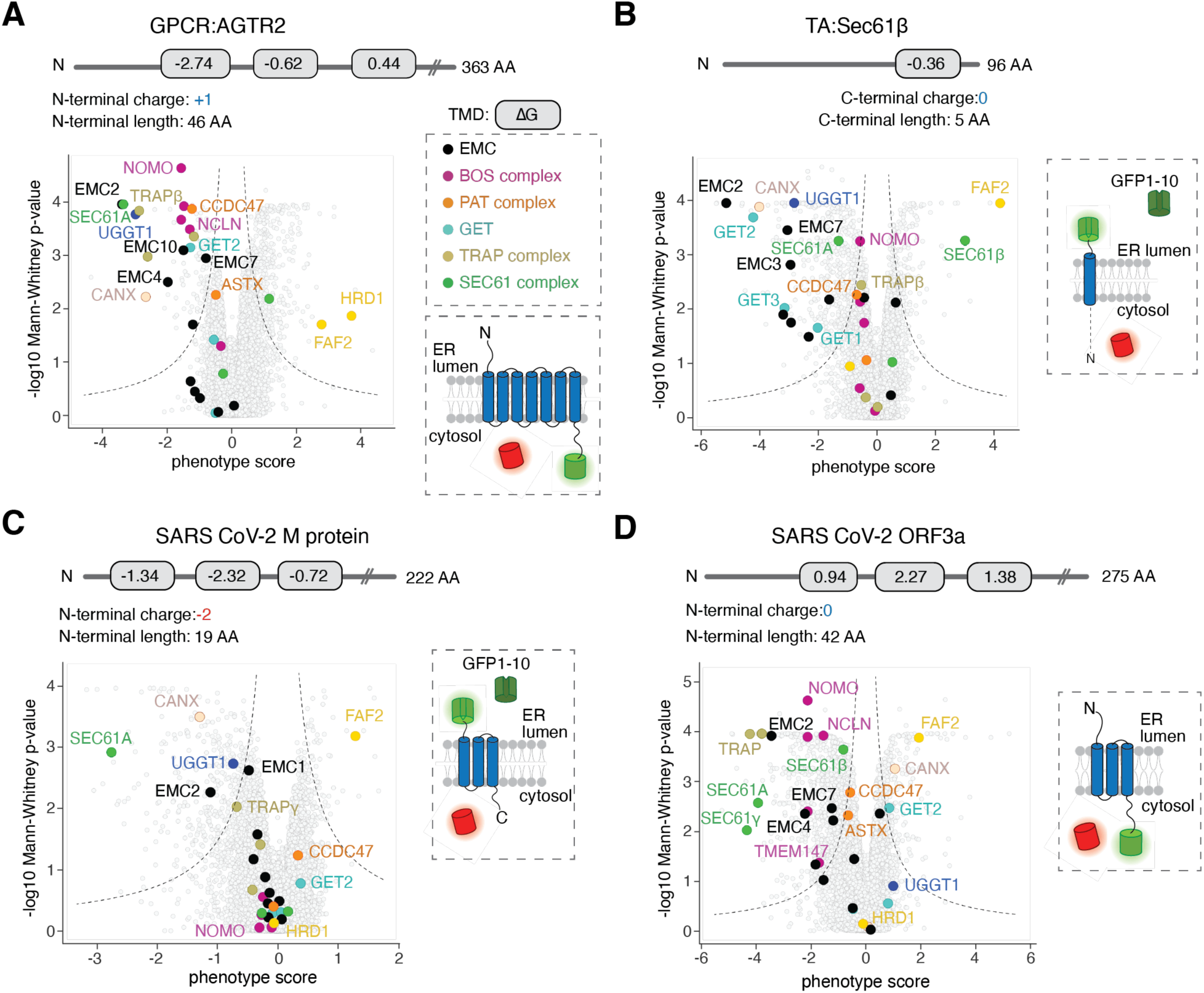
Genome-wide CRISPRi screens to systematically query biogenesis factors for diverse ER substrates. **(A)** (Top) Schematic of the GPCR reporter AGTR2. The gray rectangle indicates a transmembrane domain (TMD) with its corresponding ΔG value, calculated using the ΔG prediction server v1.0 (Hessa et al., 2007). (Right) Topology of the AGTR2 fluorescent reporter used in the genome-wide CRISPR interference (CRISPRi) screen. Full-length GFP is fused to AGTR2 and RFP is expressed as a normalization marker from the same open reading frame, separated by a P2A sequence. (Bottom) Volcano plot of the GFP:RFP phenotype [log2(High GFP:RFP/low GFP:RFP)] for the AGTR2 reporter with the three strongest sgRNAs plotted with Mann-Whitney p-values from two independent replicates. Individual genes are displayed in grey, and specific factors that increase or decrease AGTR2 stability are highlighted and labelled. Genes that fall outside the indicated dashed lines represent statistically significant hits. **(B)** As in (A), for the tail-anchored protein Sec61β (Guna, Page, et al., 2023). Here, The GFP11 sequence is appended to the C-terminal of Sec61β. Upon TA insertion into the ER, GFP11 complements with the GFP1-10 independently localized to the ER lumen, resulting in fluorescence. **(C)** As in (A) for the SARS-CoV-2 M protein. The GFP11 sequence is appended to the N-terminal of SARS-CoV-2 M with GFP1-10 expressed in the ER lumen. **(D)** As in (A) for the SARS-CoV-2 ORF3a protein. SARS-CoV-2 ORF3a is appended to full-length GFP. Abbreviations: AA, amino acids.

Second, we chose two closely related viral proteins ORF3a and M from SARS-CoV-2 (Figure 1C,D). These multipass membrane proteins both adopt an identical three-TMD topology but their TMDs have distinct biophysical properties and insertion propensities (Dolan et al., 2022; Kern et al., 2021). Further, ORF3a and M have soluble N-termini of different length (42 vs 19 amino acids) and charge (0 vs -2), which we hypothesized could alter the suite of host factors required for their biogenesis. The ability to query two topologically related proteins that are also innocuous upon overexpression was a unique advantage of using viral substrates.

Finally, as a post-translational control we included the single spanning tail anchored (TA) protein Sec61β (Figure 1B). Tail anchored proteins contain a single TMD within ∼35 amino acids of their C-terminus, and thus cannot access the co-translational biogenesis pathways typically utilized by multipass proteins (Guna, Hazu, et al., 2023; Kutay et al., 1993). The targeting and insertion of Sec61β has been extensively characterized biochemically, and therefore serves as a comparison for machinery required for biogenesis of multipass vs singlepass membrane proteins (Guna et al., 2018; Mateja et al., 2009; Stefanovic & Hegde, 2007).

Individual cell lines stably expressing these four substrates were generated in which the GFP-tagged membrane protein was expressed along with a translation normalization marker (RFP) in human K562 cells expressing the CRISPR inhibition (CRISPRi) machinery (Figure S1C) (Gilbert et al., 2014; Horlbeck et al., 2016). Previous experiments have established that when factors required for membrane protein targeting, insertion, or folding are depleted, substrates are robustly recognized and degraded by the ubiquitin proteasome pathway leading to a decrease in GFP fluorescence (Guna et al., 2018; Pleiner et al., 2020). Conversely, if membrane protein quality control is disrupted, substrates accumulate beyond normal levels, leading to an increase in GFP fluorescence. Therefore, following transduction with a genome-wide sgRNA library, cells that displayed altered substrate levels (i.e., GFP fluorescence) relative to the normalization control were sorted using florescence activated cell sorting (FACS; Figure S1C). Deep sequencing of the sgRNAs enriched in both the low and high GFP-fluorescing cells was used to identify putative biogenesis and quality control factors, respectively.

In addition to specific factors related to the unique physiologic function of each substrate (e.g., lysosomal and vesicular trafficking factors for ORF3a; Figure S1D), we identified a panel of factors that differentially affect the biogenesis or degradation of these four model substrates (Figure 1). For example, amongst the identified quality control factors, loss of the ER-localized E3 ubiquitin ligase HRD1 was found to stabilize only AGTR2, while the more general ERAD component, FAF2 was identified in all four screens (Kikkert et al., 2004; Olzmann et al., 2013). On the biogenesis side, the GET pathway components had the most pronounced effect on the tail anchored control (Sec61β), consistent with its role in post-translational insertion (Mariappan et al., 2011; Schuldiner et al., 2005, 2008; Vilardi et al., 2011). Conversely, both the translocon associated protein (TRAP) complex and the members of the multipass translocon were only significant hits for biogenesis of AGTR2 and ORF3a, but not Sec61β or M. Interestingly however, the central insertase of the multipass translocon, the GEL complex, was not a significant hit in any screen, despite near-complete depletion under these conditions (Figure S1E). The only universally identified biogenesis factor were components of the EMC, which is consistent with its established role in TA and N_exo_ TMD insertion (Chitwood et al., 2018; Guna et al., 2018).

### Distinct pathways for biogenesis and quality control of diverse substrates

To better delineate how the factors identified in the screens affected the biogenesis of a broader range of substrates, we generated a panel of 13 membrane protein reporters that represented a range of topologies and biophysical properties (Figure 2A). We included substrates with varying numbers of TMDs of distinct lengths and hydrophobicity, as well as those that differ in the structure of the intervening soluble domains. In the panel were multipass proteins in which the N-terminus must be translocated across the ER membrane (i.e., N_exo_ topology: several GPCRs, ORF3a, and M); multipass proteins in which the N-terminus will remain in the cytosol (N_cyt_ topology: TRAM2, EAAT1, GET2, and YIPF1); and single spanning (Type II: ASGR1; Type I: TRAPα); and TA proteins (SQS, VAMP2, and Sec61β). In order to allow for direct comparison, all reporters contained a full length GFP with the exception of Sec61β, whose targeting is affected by fusion with a florescent protein and therefore required use of the split GFP approach (Figure 1A) (Guna et al., 2022; Guna, Page, et al., 2023; Inglis et al., 2020).

**Figure 2.**
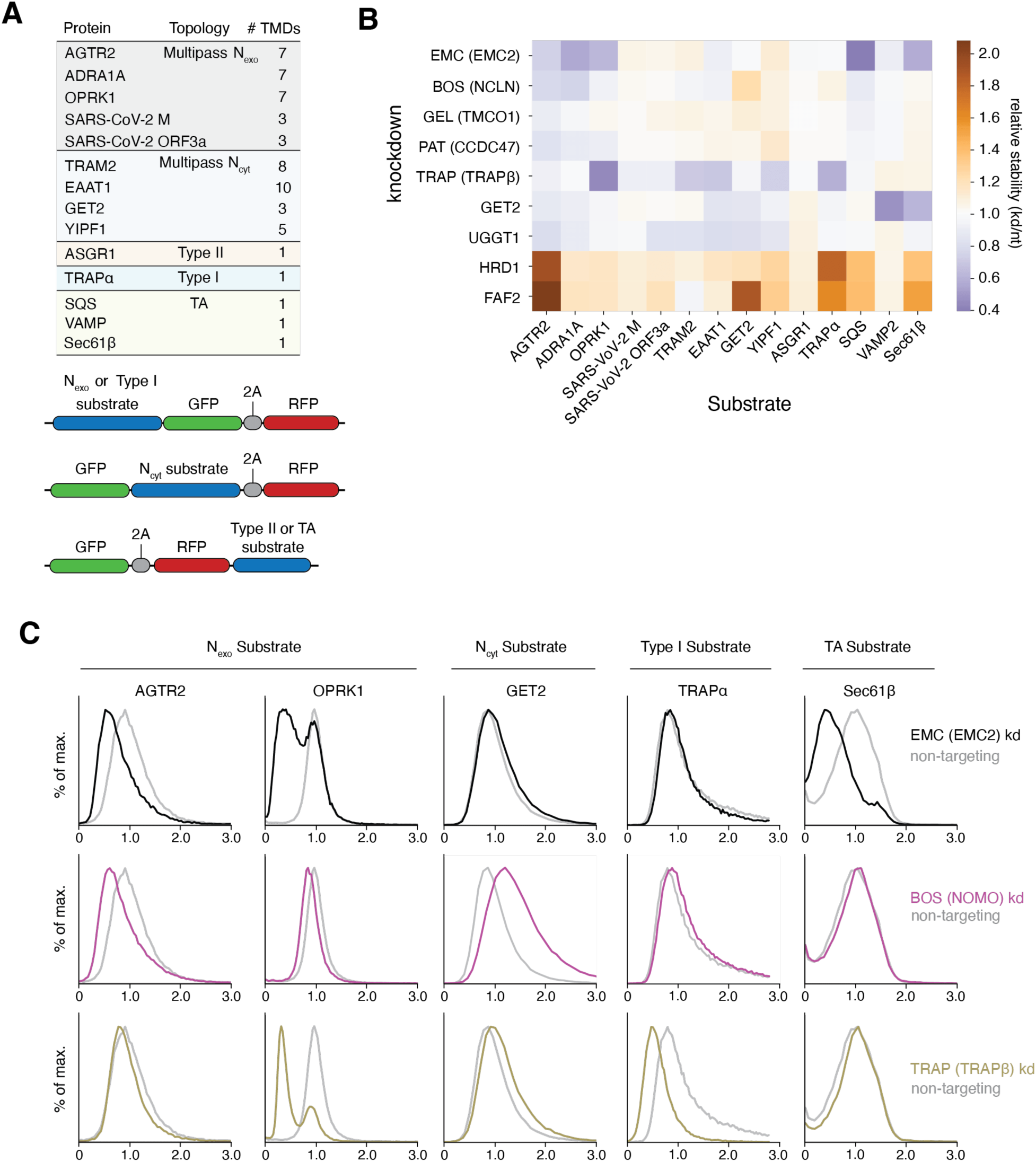
Differential effects of ER factors on membrane protein biogenesis. **(A)** (Top) Topology and number of TMDs for the panel of membrane protein reporters used to assess the effects of hits from genome-wide screens in an arrayed format. (Below) Schematic of fluorescent reporters. **(B)** Factors from forward CRISPRi screens in K562 cells (Figure 1) were systematically knocked down in RPE1 cells using CRISPRi and expression of reporters from (A) were assessed using flow cytometry. Results are shown in a heatmap indicating relative stability of each reporter after gene knockdown compared to a non-targeting control. **(C)** Representative flow cytometry analysis of individual data points from the arrayed screen summarized in (B).

Using an arrayed screen, we tested the depletion of nine factors that represented each of the major biogenesis (EMC, BOS, GEL, PAT, TRAP, GET, UGGT1) and quality control (HRD1, FAF2) complexes identified in the screens (Figure 2B). Critically, in most cases knockdown of a single subunit is sufficient to deplete the entire complex (Figure S1E) (Colombo et al., 2016; Dettmer et al., 2010; Pleiner et al., 2021; Volkmar et al., 2019). For these experiments, we used the near-diploid human RPE1 cell-line because we postulated that redundancy and compensation between factors would be more pronounced in an aneuploid cell line. Comparison across substrates both validates the hits identified in the four genome-wide screens and suggests categories of dependency that correlate with substrate topology. As expected, the clearest delineation between TAs and other membrane protein substrates is dependence on the post-translational pathway for targeting and insertion into the ER membrane. Consistent with earlier biochemical data, we find that both our forward and arrayed screens show that TA biogenesis depends on the hydrophobicity of their TMDs (Guna & Hegde, 2018; Rao et al., 2016; Shao et al., 2017; Wang et al., 2010). TAs with sufficiently hydrophobic TMDs (VAMP2) rely on the GET pathway, low hydrophobicity TAs (SQS) rely on the EMC pathway, while intermediate hydrophobicity TAs (Sec61β) can utilize both (Guna, Page, et al., 2023). Unexpectedly, loss of the GET pathway insertase GET1/2 also appeared to have a small effect on biogenesis of several multipass substrates (e.g., the GPCR AGTR2 and EAAT1) in both the genome-wide and arrayed screens.

In contrast, several factors appear to be specific to co-translationally targeted substrates. For example, depletion of the Sec61 associated chaperone, TRAP affects multipass but no TA substrates (Figure 2C) (Gemmer et al., 2023; Shao & Hegde, 2011). Though an effect of TRAPβ depletion was observed for the single spanning protein TRAPα, this may be due to an assembly rather than a biogenesis defect. While components of the multipass translocon—including the BOS, PAT, and GEL complexes—are required for substrates with multiple TMDs, they are not required for any of the single spanning membrane proteins. However, our data suggest that the function of the multipass translocon differs across cell-types, because GEL complex dependence for AGTR2 was only observed in RPE1 cells (Figure 2B) but not the K562s used in the screens, despite efficient knockdown (Figure S1E). It is possible this reflects cell-type specific changes in expression and partial redundancy and/or compensation of biogenesis factors in the ER. For example, we consistently observe that depletion of the EMC leads to a compensatory increase in TMCO1 levels (Figure S1E-F).

Critically however, even in this relatively small panel, it is clear that these multipass-specific factors are not required for the biogenesis of all multispanning proteins. For example, we observe variability in dependence on the BOS, GEL, and PAT complex amongst the three related GPCRs tested in the arrayed screen. These data suggest that biophysical properties of the TMDs and surrounding regions are more important than topology in determining biogenesis pathway. This observation sets the stage for an in-depth study of the relationship between substrate properties and biogenesis requirements.

### Identification of genetic modifiers of the EMC genome wide

One commonality across many substrates in our arrayed and genome wide screens was a dependence on the EMC for biogenesis. We therefore wondered how the EMC cooperates with other factors in the ER for insertion and folding of its multipass substrates like GPCRs. Indeed, immunoprecipitation of the human EMC from native membranes suggests it associates with a myriad of ER-resident chaperones (e.g. CNX) (Bergeron et al., 1994), biogenesis machinery (e.g. SRP receptor, glycosylation machinery, and components of the multipass translocon) (Kelleher & Gilmore, 2006; McGilvray et al., 2020; Shan & Walter, 2005), and quality control factors (e.g. the ATPase p97, responsible for extraction from the ER) (Meyer et al., 2012) (Figure 3A). Recruitment of factors required for the folding and surveillance of nascent proteins to the EMC would ensure that clients are immediately captured for maturation or degradation upon integration into the ER. These results establish the EMC as a central organizing factor for membrane protein biogenesis and quality control within the ER membrane.

**Figure 3.**
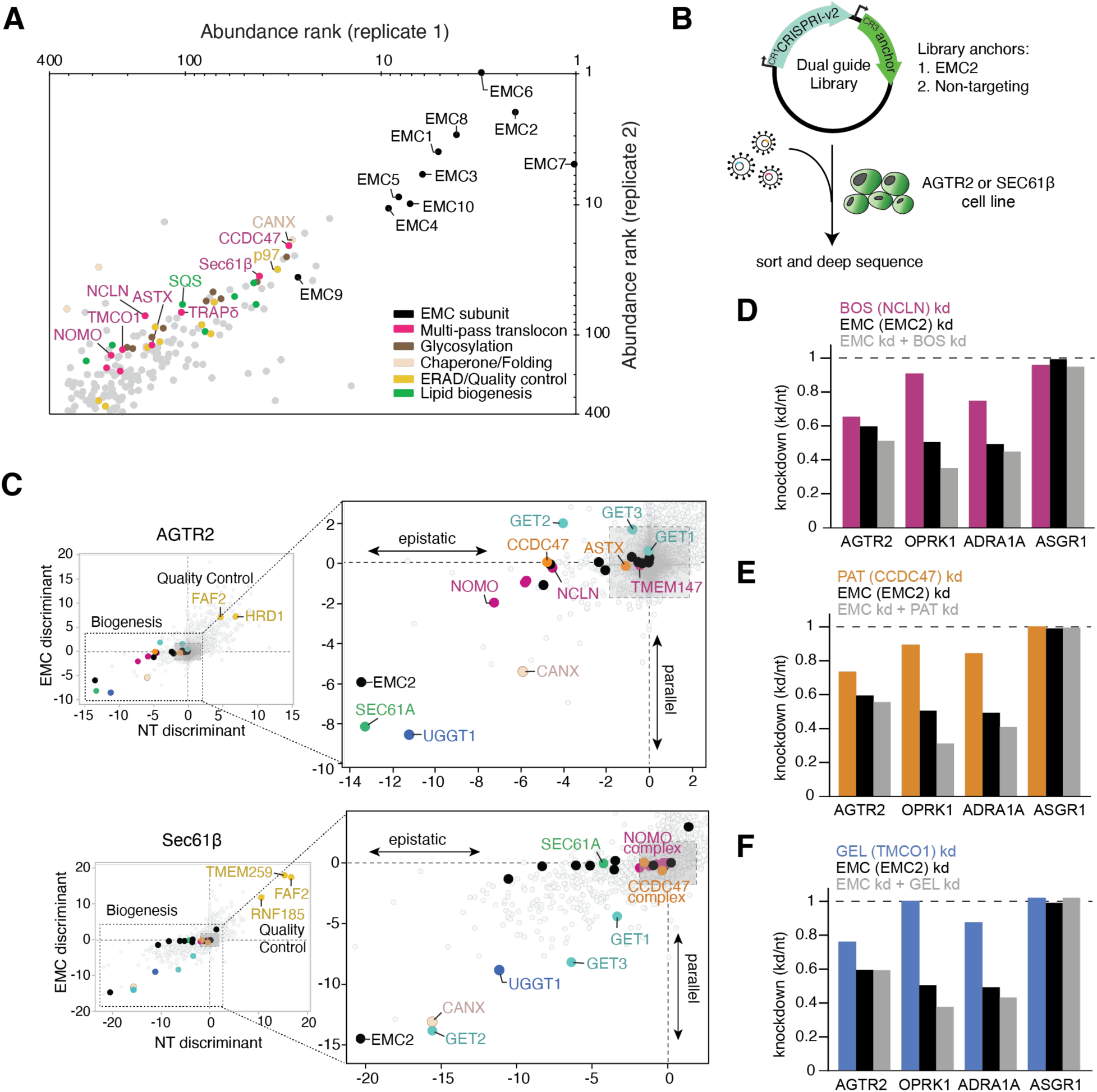
Components of the multipass translocon are epistatic with the EMC. **(A)** Scatter log-plot of abundance rank from co-immunoprecipitation of the EMC, expressed at endogenous levels, determined by mass spectrometry from two replicates. **(B)** Schematic of dual guide libraries used for genome-wide EMC genetic modifier screens in AGTR2 and Sec61β reporter cell lines. The CRISPRi-v2 sgRNA library and the genetic anchor guide, targeting either EMC2 or a non-targeting (NT) control, are expressed from a single vector (Guna, Page, et al., 2023). **(C)** Comparison of EMC2- and NT-dual library CRISPRi screens using the discriminant score as a metric (encapsulating phenotype and -log_10_ Mann-Whitney p-value for each gene) for AGTR2 (above) and Sec61β (below). Biogenesis factors are boxed and displayed at right in greater detail. Factors off-diagonal are either epistatic or parallel to the EMC, as indicated in the plots, while factors along the diagonal are independent of the EMC. **(D)** Analysis of GPCR stability (AGTR2, OPRK1, and ADRA1A) using the GFP:RFP reporter system compared to a control protein (ASGR1) as analyzed by flow cytometry after knockdown with indicated guides.

To delineate which of these associated factors are phenotypically important, we developed a dual-guide CRISPRi approach to systematically identify genetic modifiers of the EMC genome-wide (Figure 3B) (Guna, Page, et al., 2023). Briefly, we generated a library that expresses two sgRNAs on a single plasmid: (1) a genetic anchor guide, targeting the core subunit, EMC2, which when depleted results in loss of the remaining EMC subunits (Pleiner et al., 2021; Volkmar et al., 2019); and (2) a second randomized guide, targeting all open reading frames genome wide using the existing CRISPRi-v2 library (Horlbeck et al., 2016). Transduction of this dual library allows the acute knockdown of both the genetic anchor and a second randomized gene simultaneously in each cell, and is compatible with a standard CRISPRi FACS-based screening and analysis pipeline.

Comparison of the hits identified in the EMC genetic anchor screen with those from a control screen performed with a non-targeting ‘anchor’ guide library results in three categories of factors (Figure 3C and S2). First, are those that have diminished phenotypes when combined with EMC depletion, indicative of an epistatic relationship and potentially a shared pathway with the EMC. Second, are factors that have enhanced phenotypes upon loss of the EMC, likely including factors that represent parallel or partially redundant pathways. Third, are factors that act independently of the EMC, and therefore show no change in phenotype with or without the EMC.

Interrogating both the EMC-dependent tail-anchored protein Sec61β and the GPCR AGTR2 allowed us to delineate EMC co-factors that function to support its post vs co-translational biogenesis roles (Figure 3C and S2). Validating this approach, all EMC subunits have a diminished effect in the EMC2 knockdown background for both TA and GPCR biogenesis. Conversely, the phenotype of known parallel pathways for TA insertion, including the GET components, are enhanced by EMC depletion, particularly in the TA screen (Figure S2C,D). Finally, several quality control factors, such as HRD1 and FAF2 exhibit EMC independent effects, suggesting their function may be agnostic to the insertion pathway.

Direct comparison of the biogenesis factors identified in the TA vs the GPCR genetic modifier screens suggest that many more factors are cooperating with the EMC in insertion and folding of multipass membrane proteins than of TAs. In particular, we identified many components of the multipass translocon, including subunits of the BOS and PAT complexes as epistatic with the EMC for AGTR2 biogenesis (Figure 3C-F and S2A,B). To test whether other GPCRs display a similar epistatic dependence, we performed an arrayed screen with dual guides targeting the BOS, PAT, or GEL complexes alone or in combination with a guide targeting the EMC. We included the GPCRs AGTR2, OPRK1 and ADRA1A and the type II membrane protein ASGR1 (Figure 4D-F). For the GPCRs but not for ASGR1, EMC displays an epistatic relationship with the BOS, PAT, and GEL complexes, suggesting that this relationship may be a general feature of multipass membrane protein biogenesis. The BOS, PAT, and GEL complexes were further found to co-purify with the EMC under conditions where all components were expressed at endogenous levels (Figure 4A-B), suggesting both a genetic and physical interaction between these biogenesis machineries.

**Figure 4.**
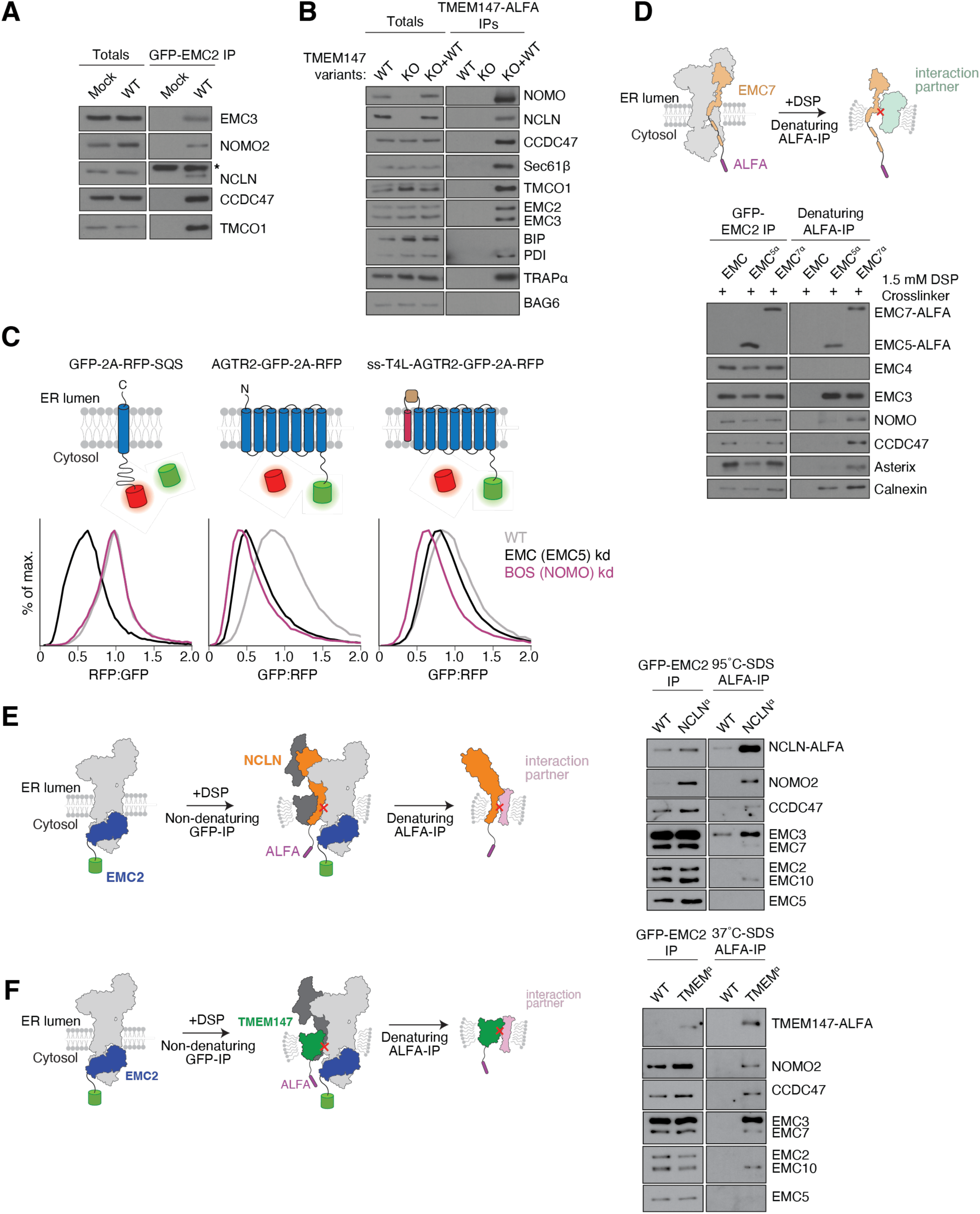
The EMC and the BOS complex are direct physical interactors. **(A)** Cell lines stably expressing GFP-EMC2 were solubilized in the detergent GDN, immunoprecipitated under native conditions using anti-GFP nanobody, and analyzed by SDS-PAGE and western blotting compared to a mock control. An asterisk indicates a cross-reacting band. **(B)** As in (A) with anti-ALFA nanobody immunoprecipitation of TMEM147-ALFA expressed at endogenous levels in a TMEM147 KO cell line. **(C)** Flow cytometry analysis of the ratiometric SQS and AGTR2 protein reporters described Figure 2. The signal sequence (ss) of Pre-Prolactin followed by T4 Lysozyme (T4L) was appended to the N-terminal of AGTR2. RPE1 cells were treated with siRNAs targeting EMC5, NOMO, or a non-targeting control (WT) and transduced with the appropriate fluorescent reporter construct. Histograms of the RFP:GFP ratio (SEC61β) or GFP:RFP ratio (AGTR2 with and without the signal sequence) are shown. **(D)** Chemical crosslinking and immunoprecipitations from stable cell lines expressing GFP-EMC2 alone, GFP-EMC2 and EMC7-ALFA or GFP-EMC2 and EMC5-ALFA. The crosslinker DSP is thiol reducible, and the samples were analyzed using SDS-PAGE under reducing conditions and western blotting. **(E)** As in (D), but for cell lines stably expressing GFP-EMC2 and NCLN-ALFA. **(F)** As in (D) for cell lines expressing GFP-EMC2 and TMEM147-ALFA. Note that since TMEM147 almost quantitatively precipitates after boiling in SDS, denaturation at 37°C was chosen to enable its immunoprecipitation.

One potential trivial explanation for genetic epistasis between the EMC and BOS complex is that, following insertion of TMD1 of a GPRC by the EMC, the multipass translocon is responsible for inserting the remaining downstream TMDs. However, we found that addition of a signal sequence or signal anchor to the N-terminus of AGTR2 or the GPCR ADRA1A, which allows them to bypass the EMC and utilize Sec61 for insertion of its first TMD (Chitwood et al., 2018), markedly rescues its dependence on the BOS complex for biogenesis (Figure 4C and S3C-D). Indeed, the rescue of BOS complex dependence upon addition of a signal sequence is a similar magnitude to that observed for the EMC. We therefore concluded that there may be an additional role of the BOS complex at the EMC, beyond its previously reported function at as part of the multipass translocon. We therefore sought to determine whether the BOS complex may function as a co-factor of the EMC in biogenesis of multipass membrane proteins.

### The BOS complex is a direct physical interactor of the EMC

Though the BOS and PAT complexes co-immunoprecipitated with the EMC, we first sought to confirm that this reflected a direct physical interaction. To do this we incubated intact cells under conditions where the EMC (Figure S3A) and the BOS complex (Figure S3B) components are present at endogenous levels with the amine-reactive chemical crosslinker DSP. DSP has a length of ∼12Å, such that only factors within close proximity can be covalently crosslinked. The resulting crosslinked species were immunoprecipitated under denaturing conditions in SDS, where we found that subunits of the BOS (NOMO) and PAT (CCDC47) complexes specifically immunoprecipitated with EMC7 under conditions in which other EMC subunits are markedly depleted (Figure 4D). NOMO did not crosslink to EMC5, confirming that the limited crosslinking conditions, and thereby co-immunoprecipitation with EMC7, was specific.

To confirm that this result did not reflect a long-range interaction between the flexible IgG domains of NOMO and the EMC, we performed similar experiments using affinity tags on the other BOS complex subunits that are either more rigid (NCLN) or fully embedded in the bilayer (TMEM147). We found that immunoprecipitation of TMEM147 and NCLN after chemical crosslinking specifically recovered EMC3,7, and 10, but not EMC2 (Figure 4E-F). We therefore concluded that the BOS complex is a direct physical interactor of the EMC in native membranes.

To better understand the interaction between the EMC and the BOS complex, we sought to determine a structure of the 12-subunit holocomplex purified from human cells. Though we have shown that the BOS complex and the EMC interact without exogenous stabilization (Figure 4A-B), to increase their local concentration and thereby enable structural analysis, we introduced an ∼50 amino acid linker between TMEM147 and EMC2. By using an extremely long and flexible linker, we avoid artificially stabilizing a non-physiologic interaction between the EMC and BOS complex. Indeed, modeling suggested that this >100 Å linker would not preclude interaction of the EMC and BOS complexes in any orientation or arrangement. Using single particle cryo-EM, we determined a modest resolution structure of the EMC•BOS holocomplex to an overall resolution of 5.2 Å in the detergent GDN (Figure 5A and S4A). From the EM density, it is clear that BOS and the EMC are within the same detergent micelle, and interact at a fixed orientation relative to each other.

**Figure 5.**
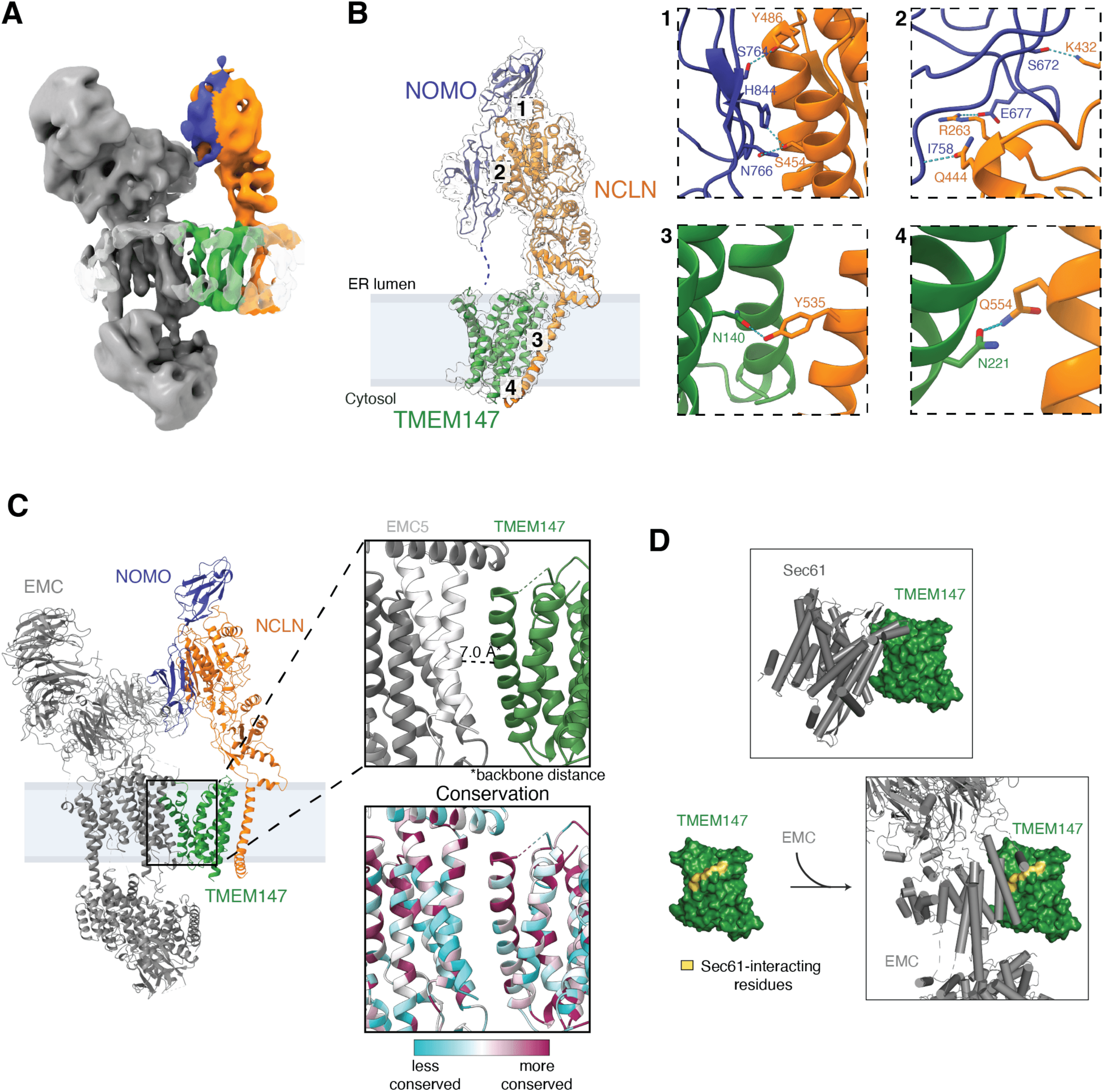
Structural analysis of the EMC•BOS holocomplex. **(A)** Coulomb potential map of the EMC•BOS complex. **(B)** In order to enable de novo modeling of the BOS complex, we isolated two different versions containing either full length NOMO with all 12 IgG repeats (fNOMO), or a truncated version including only the 3 c-terminal IgG domains (tNOMO). The molecular model of the BOS(tNOMO) complex (BOS complex with truncated NOMO) with its corresponding EM density map is shown. Insets 1-4 show magnified regions where the subunits are in close contact. Contacting residues are labeled accordingly. Insets 1 and 2 show contacts in the lumen, while 3 and 4 show contacts in the membrane. **(C)** Molecular model of the BOS (fNOMO) • EMC complex. The top right image shows a close-up view between EMC5 and TMEM147 to illustrate their proximity, with the backbone distance being 7.0 Å in one instance. The bottom right image shows the conservation of the same region, with the less conserved residues colored in cyan and more conserved residues colored in dark magenta. **(D)** Comparison of the interacting surfaces with TMEM147 of Sec61 and EMC. Top: Cylinder cartoon representation of Sec61 interacting with TMEM147 shown in surface representation (PDB 6W6L) (McGilvray et al., 2020). Bottom left, the TMEM147 residues that are interacting with Sec61 are highlighted in yellow. These residues were determined using the *InterfaceResidues.py* script from the PyMOL wiki (https://pymolwiki.org/index.php/InterfaceResidues). Right: Cylinder cartoon representation of EMC interacting with TMEM147 with the Sec61-interacting residues being highlighted to show the overlap of interacting surfaces between Sec61 and EMC.

While we could unambiguously fit existing models of the isolated EMC into the holocomplex density map, the resolution was insufficient for de novo building of the BOS complex. Therefore, using an affinity tag on TMEM147 and more stringent conditions, we purified the isolated BOS complex and determined two structures: (1) BOS(fNOMO), using the full-length NOMO, including its 12 endogenous IgG domains (∼8 Å resolution) (Figure S4B and S5A); and (2) BOS(tNOMO) in which we truncated all but the last 3 IgG repeats of NOMO (3.7 Å resolution) (Figure 5B and S4C). Notably, the truncated NOMO resulted in a higher purity protein sample and improved monodispersity upon freezing on an EM grid. In both structures, only the 2-3 terminal IgG domains of NOMO interact with the lumenal domain of NCLN, suggesting that the remaining IgG repeats are likely dynamic. Indeed, the previously reported structure of the multipass translocon complex (McGilvray et al., 2020; Smalinskaitė et al., 2022) could not unambiguously assign density for any of the IgG domains. Superposition of the density of the full-length NOMO and truncated NOMO in both complexes suggest that they are qualitatively identical, validating use of the truncated complex for high-resolution model building (Figure S5A).

We then used this model to unambiguously fit the EM density in the EMC•BOS holocomplex. The EMC and BOS interaction forms a buried surface area of ∼855 Å^2^. There are two primary interfaces: both an intramembrane interface, composed primarily of EMC5 and TMEM147, as well as a smaller lumenal interface between NCLN and EMC1. Notably, the intermembrane surface between the EMC and BOS complexes is composed primarily of conserved hydrophobic residues in EMC5 and TMEM147 (Figure 5C and S5C), suggesting that this interface may be the more important than that in the lumen. We hypothesize that the absence of crosslinks previously observed between EMC5 and NOMO (Figure 4D) can be explained by the lack of primary amines in the membrane-embedded EMC5 subunit. Finally, the interaction surface with the BOS complex is distinct compared to that reported for the chaperone-binding mode of EMC (Figure S5B) (Chen et al., 2023).

Comparison with structures of the multipass translocon bound at Sec61 suggest that BOS binding to the EMC and Sec61 is mutually exclusive (Figure 5D) (McGilvray et al., 2020). Conversely, binding of BOS to the GEL and PAT complexes would all be compatible with interaction at the EMC (Figure S5D). Indeed, we observed co-immunoprecipitation of both PAT and GEL complex subunits with the EMC by both quantitative proteomics and western blotting, and observed that their interaction with the EMC is independent of EMC-BOS interaction (Figure 3A,4A and S5E). Further, we verified using chemical crosslinking that CCDC47 is a direct physical interactor of the EMC and that its interaction is EMC7-dependent (Figure 4D, S5F). EMC7 is a peripheral subunit of the EMC, and as such, knockout of EMC7 does not destabilize core EMC subunits (Pleiner et al., 2023). This therefore suggests that the interaction between the EMC and CCDC47 is highly specific, as loss of a single peripheral subunit abolishes CCDC47 interaction with the EMC. Cumulatively, these data suggest that the multipass translocon, including the BOS, PAT, and GEL complexes, are also assembled at the EMC in a mutually exclusive manner to their binding to Sec61.

### Biophysical properties of substrate soluble domains dictate biogenesis pathway

Having established both a genetic and physical interaction between the EMC and BOS complex, we sought to determine the function of the BOS complex in the biogenesis of EMC-dependent substrates. Analysis of the genome-wide and arrayed screens revealed patterns in the biophysical and topological features that confer dependence on the BOS complex during biogenesis. First, it was clear that the function of the BOS complex is specific to multipass membrane proteins, as it was not identified in the TA screen, and its depletion did not affect single spanning substrates in our arrayed panel (Figure 1,2). However, not all multipass substrates are equally dependent on the BOS complex. We therefore reasoned that if the EMC is specifically required for insertion of the first TMD of these multipass substrates, dependence on the BOS complex may be conferred by properties of this TMD and its surrounding sequence.

Comparison of the M and ORF3a screens suggested that one marked difference between these two topologically identical proteins, was the length and charge of the N-terminus of ORF3a (Figure 1C,D). Indeed, AGTR2 and ADRA1A, which also displayed a clear reliance on the BOS complex when utilizing the EMC (Figure 4C and S3C,D), contain an atypically positively charged soluble N-terminus in comparison to most GPCRs that lack signal sequences (Wallin & von Heijne, 1995). Previous experiments found that the EMC uses a positively charged hydrophilic vestibule for insertion, which limits integration of substrates containing positively charged soluble domains (Pleiner et al., 2023; Wu & Hegde, 2023). Consistent with this observation, most GPCRs and ER-targeted TA proteins contain neutral or negatively charged soluble domains that would be translocated through the hydrophilic vestibule of the EMC. We therefore hypothesized that when the first TMD of a multipass protein is inserted by the EMC, the charge of the N-terminal soluble domain confers dependence on the BOS complex.

To test this, we selected a suite of GPCRs and tested their dependence on EMC, BOS, and PAT complexes for biogenesis (Figure 6A and S6A,B). We found that in general, those substrates with more positively charged N-termini displayed a stronger dependence on the BOS complex (Figure 6A), but observed no connection between charge and EMC dependence (Figure S6A). However, this correlation was imperfect, suggesting that features beyond simply charge may play an important role.

**Figure 6.**
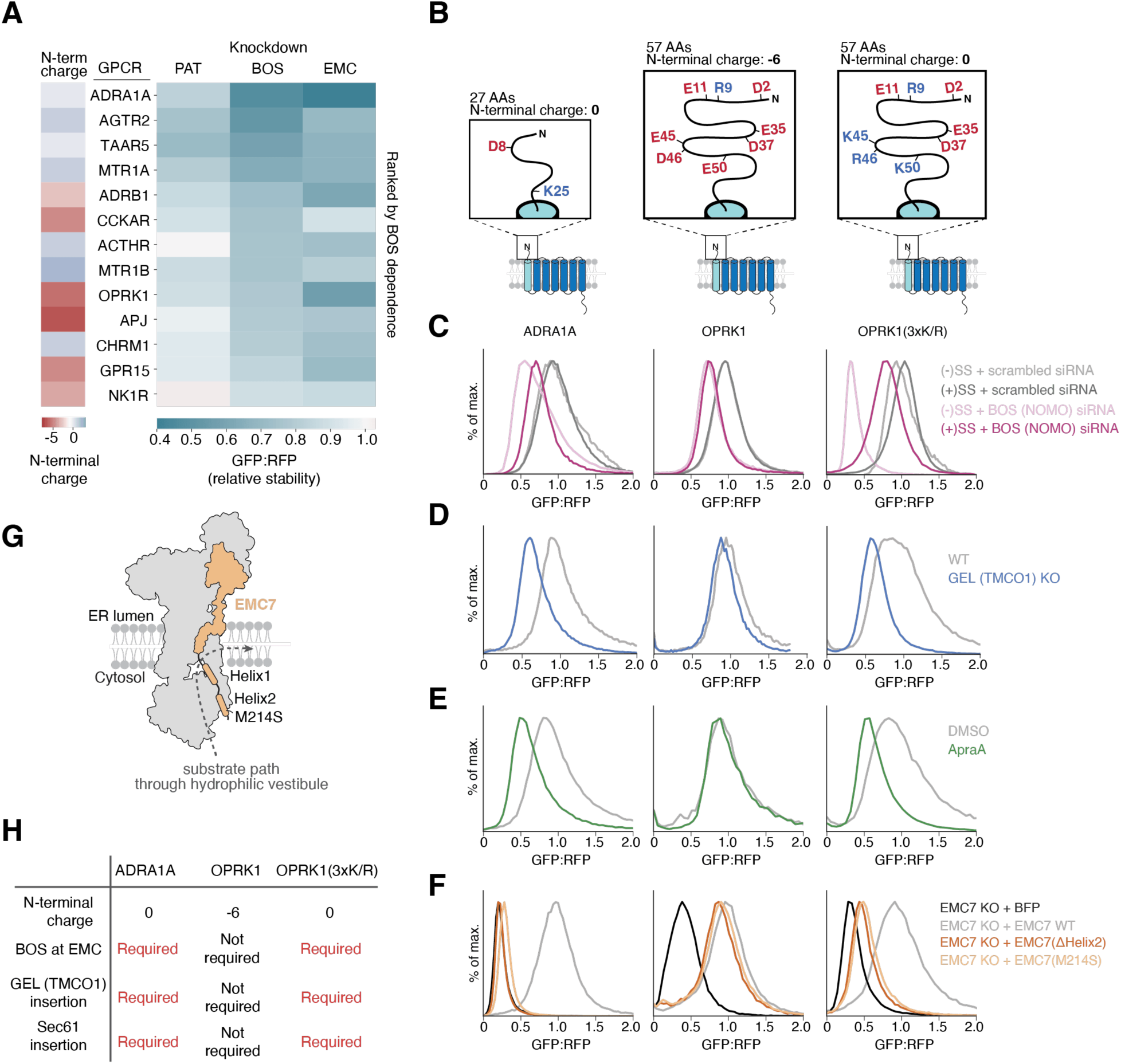
Charge of the N-terminal domain of EMC substrates determines the requirement of the BOS complex at EMC. **(A)** The indicated GPCRs were expressed in the GFP-2A-RFP cassette after siRNA knockdown of EMC5, NOMO, CCDC47, or a non-targeting control in RPE1 cells and analyzed by flow cytometry. The relative stability of each GPCR was determined as a ratio of GFP:RFP in the knockdown condition compared to the control. (Right) Shown is a heatmap of the relative stability of each GPCR, ranked by BOS dependence. (Left) A heatmap indicating the charge of the N-terminal domain of each GPCR. **(B)** Schematics of GPCRs ADRA1A, OPRK1 and a variant of OPRK1 with a net neutral N-terminal domain (3xKR). **(C)** Histograms from flow cytometry assays with ADRA1A, OPRK1, or OPRK1(3xKR) with or without the Pre-Prolactin signal sequence (SS) and T4L Lysozyme, as described in Figure 4C. Cells were treated with siRNAs targeting NOMO or a scrambled control sequence. **(D)** As in (C), in WT or TMCO1 KO cells and with the signal anchored GPCR reporters. **(E)** Reporters in (D) were assayed for SEC61-dependent insertion using the inhibitor Apratoxin A. Cells were analyzed as in (D). **(F)** Reporters in (D) were assayed for EMC7 dependence using flow cytometry. EMC7 KO cells were transduced with either a BFP control, WT EMC7, EMC7 with a deletion of Helix2 (ΔHelix2), or EMC7 containing a single point mutation in Helix2 (M214S) before analysis as described in (D). **(G)** Cartoon of the EMC with EMC7 highlighted showing a substrate’s path of insertion into the ER bilayer (Pleiner et al., 2023). **(H)** Summary of data in (C)-(F).

To interrogate the relationship between N-terminal charge and BOS complex dependence directly, we chose two representative GPCRs, OPRK1 and ADRA1A. Both substrates display strong EMC dependence, but contain distinct N-terminal charge (−6 vs +0) (Figure 6B). As previously observed for AGTR2, ADRA1A only requires the BOS complex when utilizing the EMC for insertion of its first TMD (Figure 4C, 6C and S6C,D). In contrast, while addition of a signal sequence to the N-terminus of OPRK1 rescues the effect of EMC depletion, it has minimal effect on the BOS complex dependence. However, an OPRK1 mutant with three additional positive charges within its N-terminus no longer relies on the BOS complex when inserted by Sec61. We therefore concluded that by studying the insertion of these three substrates we could precisely test the effects of the soluble N-terminus of a GPCR on its biogenesis.

Given earlier data suggesting that positively charged soluble domains are less efficiently translocated by the EMC during insertion, we wondered if these substrates might be more likely to rely on alternative insertase pathways. To do this, we tested whether GPCRs displayed differential dependence on the GEL complex (i.e., TMCO1) and the insertase activity of Sec61. To prevent pleotropic effects of Sec61 depletion and specifically query the role of its insertase activity, we used the inhibitor Apratoxin A, which prevents opening of the Sec61 lateral gate (Paatero et al., 2016; Thornburg et al., 2013). We found that substrates containing a positively charged N-terminus had increased dependence on TMCO1 and the lateral gate of Sec61. Importantly, while wild type OPRK1 displayed little or no dependence on TMCO1 or Sec61, the positively charged N-terminal OPRK1 mutant showed increased dependence on both alternative insertases (Figure 6D,E and S6D). Moreover, we observed an even greater dependence of the positively charged N-terminal OPRK1 mutant on TMCO1 and Sec61 when both insertases were impaired simultaneously, both in cells and in vitro (Figure S7). We therefore concluded that those multipass substrates that cannot be efficiently inserted by the EMC due to charge in their soluble domains instead rely on alternative, partially redundant, pathways for biogenesis.

Finally, we tested if these substrates might have increased dependence on the flexible, methionine-rich loops of the EMC that were previously shown to physically interact with substrates in the cytosol during their passage into the membrane (Pleiner et al., 2020, 2023). Indeed, GPCRs containing a positively charged N-terminus display increased dependence on the C-terminus of EMC7 that directly binds substrate TMDs below the hydrophilic vestibule of the EMC (Figure 6F,G and S6F). Comparison of the wild type and N-terminal positively charged mutant OPRK1 suggest that N-terminal charge is sufficient to increase the dependence on these flexible cytosolic domains. An increased reliance on these cytosolic domains of the EMC would be consistent with an increased dwell time at the EMC for positively charged substrates that are not efficiently translocated through the positively charged vestibule. This could also provide a potential mechanism by which substrates are transferred to alternative insertases (i.e., TMCO1, Sec61) if not immediately inserted by the EMC.

## DISCUSSION

### Topology and biophysical properties dictate biogenesis pathway

For many decades it was postulated that the Sec61 translocation channel provided the major pathway into the ER for integral membrane proteins (Shao & Hegde, 2011). However, it is clear that Sec61 alone is insufficient for the insertion and folding of most ER-targeted substrates. By systematically studying the biogenesis of membrane proteins using a series of forward and arrayed genetic screens, we have begun to dissect the substrate specificity of the suite of biogenesis factors in the ER (Figure 1,2). Multiple insertases beyond Sec61, including the Oxa1 superfamily members GET1/2, EMC, and the GEL complex, are required for insertion of many single spanning and multipass membrane proteins. As expected, the post-translational insertase GET1/2 had the most pronounced role in TA biogenesis, but its depletion also affected some multipass substrates. This result could be indirect, resulting from increased flux of endogenous TAs through competing pathways in the absence of GET1/2. However, it is also possible that under some circumstances, GET1/2 can play a broader role in membrane protein biogenesis at the ER. In contrast, loss of the GEL complex, the insertase component of the multipass translocon, had no effect on single-spanning membrane proteins.

However even amongst proteins of identical topology, we observed distinct biogenesis requirements. For example, comparison of several GPCRs showed that the Sec61-associated chaperone TRAP, was required for OPRK1 biogenesis but not ADRA1A (Figure 2). Conversely, loss of the BOS complex had a more pronounced effect on ADRA1A and AGTR2 than OPRK1. Therefore, our data suggests that both topology and the biophysical properties of the substrate must determine the biogenesis pathway.

Earlier work has established that the hydrophobicity of a substrate TMD dictates biogenesis requirements, in some cases requiring unique insertion machinery (i.e. for TAs) or chaperones (i.e. Asterix) (Chitwood & Hegde, 2020; Guna, Hazu, et al., 2023; Meacock et al., 2002). Our data suggests that it is not only the TMD that can confer dependence on particular biogenesis pathways, but also the biophysical properties of the associated soluble domains. For example, while wild type OPRK1 shows little to no dependence on the GEL complex or Sec61 for insertion of its first TMD, a mutant that contains three point mutations within the soluble N-terminus confers dependence on both alternative insertases. The immediate consequence of these observations is that sequences distant from the TMD, in some cases as far as 50 amino acids away, have a profound effect on both insertion efficiency and biogenesis pathway into the ER. It is therefore clear that not only are the biophysical properties of the TMD itself, but also its associated soluble domains an important consideration for biogenesis of diverse membrane proteins.

### The EMC is a central organizing hub for biogenesis and quality control

Having delineated roles for individual factors in the ER, we used a series of functional and structural approaches to define how these components cooperate to achieve insertion and folding of diverse membrane proteins. Comparison of the forward and arrayed screens place the EMC at the center of the membrane protein biogenesis and quality control machinery in the ER. Biochemically, we observed tight coupling of the EMC with quality control machinery (e.g., p97), which ensures mislocalized or mis-inserted membrane proteins are efficiently triaged for extraction and degradation (Figure 3A). Conversely, co-localization of the EMC with ER resident chaperones (e.g., CANX) and post-translational modification machinery (e.g., glycosylation) guarantees that nascent substrates are immediately captured for modification and folding upon integration into the ER. This type of local recruitment of biogenesis and quality control machinery to the site of protein insertion is analogous to that observed for the translocation channel, Sec61 (Gemmer et al., 2023; Görlich et al., 1992; Hartmann et al., 1993; Jaskolowski et al., 2023; Kalies et al., 1998; Voigt et al., 1996). Further, direct recruitment of the multipass translocon components, including the BOS, GEL, and PAT complexes to the EMC facilitates organization of ‘biogenesis hubs’ in the ER membrane. The observation that at least some of these factors are stabilized by interaction with EMC7 (Figure S5F) provides one potential explanation for the conservation of the large soluble lumenal and cytoplasmic domains of the EMC: recruitment and retention of auxiliary factors to the site of membrane integration.

### A working model for multipass membrane protein insertion

The organization of biosynthesis machinery into local and dynamic hubs within the ER— including the EMC, Sec61, and the multipass translocon components—provides a coherent working model for the insertion and folding of complex multipass membrane proteins. A ribosome nascent chain complex is delivered to the ER by the signal recognition particle (SRP), where substrates first probe the hydrophilic groove of the EMC. Though the molecular details of handover from SRP to the EMC are not yet precisely defined, SRP receptor subunits were recovered in native co-immunoprecipitation of the EMC from ER membranes, suggesting one potential mechanism for recruitment to the EMC (Figure 3A). Models consistent with a ‘first refusal’ of substrates by the EMC, best explain data showing the EMC can enforce the correct folding of multipass substrates containing positively charged extracellular domains (Pleiner et al., 2023). Our observation that the PAT complex directly interacts with the EMC, including CCDC47, which is known to use its soluble coiled-coil domain to bind the ribosome, provides a putative mechanism for transiently stabilizing ribosome nascent chain complexes at the EMC (McGilvray et al., 2020; Smalinskaitė et al., 2022; Sundaram et al., 2022). Recruitment of the PAT complex in an EMC7-dependent manner also provides one potential explanation for the differential dependence of substrates on EMC7 (Figure 6F).

Substrates in which the first TMD can be efficiently inserted by the EMC, such as those with negatively charged and short N-terminal soluble domains, passage through its hydrophilic vestibule into the bilayer. Co-localization of the EMC with glycosylation machinery would then allow immediate post-translational modification of the soluble N-terminus. In contrast, substrates that are poorly inserted by the EMC (i.e., those that have increased N-terminal positive charge but also likely including those with longer or more structured soluble domains) have a longer dwell time at the cytosolic vestibule of the EMC. This is consistent with their increased reliance on the cytosolic C-terminus of EMC7 that contains several conserved hydrophobic residues, previously shown to directly interact with substrates in the cytosol (Pleiner et al., 2023). These data are consistent with the model that the rate limiting step for insertion is translocation of the N-terminal soluble domain through the hydrophilic vestibule of the EMC. The properties of the N-terminus therefore dictate the energetic barrier for translocation into the ER lumen.

TMDs that are not immediately inserted by the EMC are shuttled to alternative insertases including the GEL complex (i.e., TMCO1) and in some cases Sec61. We hypothesize that handover of those substrates with charged N-terminal domains between insertases is facilitated by recruitment of the BOS, GEL, and PAT complexes to the EMC. Indeed, these substrates appear to primarily require the activity of the multipass translocon when using the EMC for insertion of their first TMDs (Figure 6C).

Structures of the EMC•BOS holocomplex are most consistent with a model in which interactions between biogenesis machineries are dynamic. Rather than a rigid holocomplex that includes both the EMC and Sec61, co-localization may serve primarily to increase local concentration of insertases to accommodate diverse substrates as they passage into the bilayer. This would be consistent with the observation that recruitment of the BOS complex to the EMC is mutually exclusive to its binding to Sec61 (Figure 5D) (McGilvray et al., 2020; Smalinskaitė et al., 2022). Mutually exclusive binding of the multipass translocon components to the EMC may also encourage handover to Sec61. Certainly, recruitment of the multipass translocon components to the EMC provides a putative mechanism for how any substrate in which the first TMD is inserted by the EMC is transferred to the back of Sec61 to complete its insertion and folding.

### A central role for Oxa1 superfamily of insertases throughout all kingdoms of life

Based on differences in their biophysical properties (e.g., charge of the hydrophilic vestibule, limitations on length/charge/structure to be translocated), it is likely that insertases display partial substrate preferences (Figure 7A). However, it is clear that Oxa1 superfamily insertases such as the EMC and GEL complex are partially redundant, and loss of any one results in compensation in human cells (Figure S1F). Remarkably, recruitment of the core insertase subunits EMC3/6 to the inner mitochondrial membrane was sufficient to rescue loss of Oxa1 in yeast (Güngör et al., 2022). Further, the ability of one Oxa1 superfamily insertase to compensate for loss of another explains genetic studies that suggest loss of function mutations in TMCO1 are not deleterious in the human population (Karczewski et al., 2020). In these cases, upregulation of the EMC or potentially even GET1/2 may be sufficient to ensure efficient insertion of all required membrane protein substrates.

**Figure 7.**
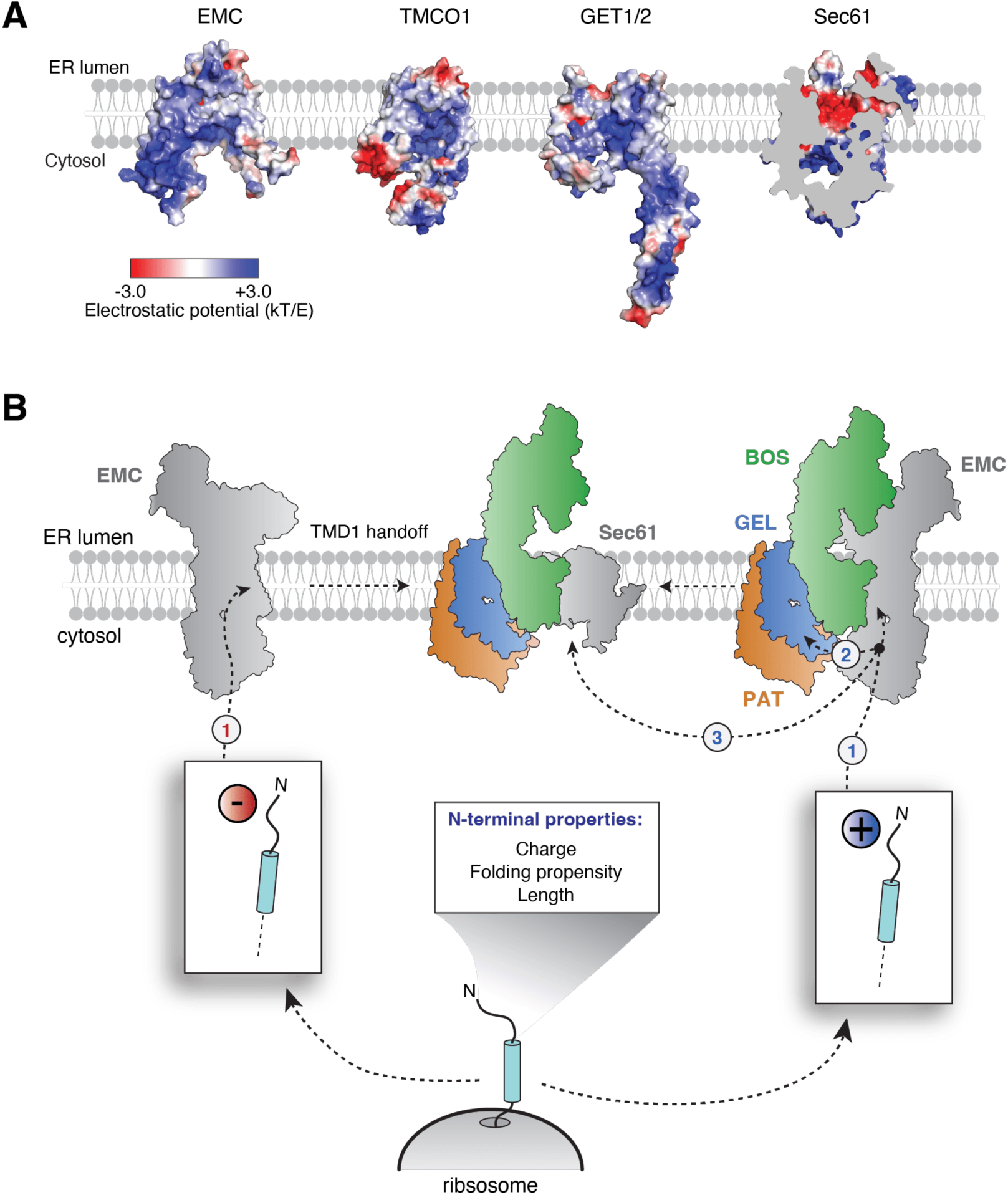
Model of multipass membrane protein biogenesis at the ER. **(A)** Comparison of the electrostatic surfaces of known human insertases at the ER membrane. Electrostatic potential was generated using APBS with range -3 to +3 kT/e and mapped onto surface representation of EMC3/6 (PDB ID 8S9S) (Pleiner et al., 2023), TMCO1 (PDB ID 6W6L) (McGilvray et al., 2020), GET1/2 (PDB ID 6SO5) (McDowell et al., 2020), and Sec61 (PDB ID 3JC2). (Voorhees & Hegde, 2016) **(B)** Model for membrane protein insertion at the ER. Multipass N_exo_ proteins are co-translationally targeted to the EMC, where the properties of their N-terminal soluble domain alter the requirements for insertion of the first TMD. (Left) Proteins with a net negative charge in their N-terminal domain are able to utilize the EMC for TMD1 insertion without additional co-factors. After insertion of TMD1, these proteins are then handed off to the multipass translocon at Sec61 for insertion of the remaining TMDs. (Right) Proteins that have net neutral or positive N-terminal domains require additional factors for insertion of the first TMD. The BOS complex physically associates with the EMC and is required for EMC-dependent insertion of the first TMD. Additionally, these substrates, are routed to TMCO1 or Sec61 for TMD1 insertion. After TMD1 is inserted, the multipass translocon at Sec61 inserts the remaining TMDs. Binding of the BOS, GEL, and PAT complex to the EMC may facilitate hand-over of substrates between the EMC and the multipass translocon during membrane protein biogenesis. This co-translational triaging of ribosome nascent chain complexes with exposed TMD1 based on N-terminal domain features is likely dictated by the biophysical properties and architecture of the EMC’s hydrophilic vestibule. Charge repulsion between EMC’s positively charged hydrophilic vestibule and its clients’ N-terminal domains would increase dwell time and facilitates the engagement of multipass translocon factors and insertion via TMCO1 or Sec61.

This redundancy between Oxa1 superfamily insertases explains how membrane protein biogenesis is achieved across evolutionarily distinct kingdoms of life (Anghel et al., 2017; Kumazaki et al., 2014). Superficially, it appears that the multipass translocon components are a metazoan specific adaptation, without homologs in yeast and bacteria. For example, while the EMC is conserved in all eukaryotes, the GEL complex is metazoan specific (Anghel et al., 2017). However, it is clear that even in humans, while the EMC is essential, the presence of both the EMC and GEL complexes is not strictly required. Interestingly, in fungi, which may rely primarily on the EMC for multipass insertion, the EMC is missing one of its soluble subunits (i.e., EMC8/9) (Jonikas et al., 2009). The lack of EMC8/9 would allow the EMC to sit closer to the ribosome, potentially taking the place of the GEL complex in the multipass translocon. An analogous reliance on Oxa1 superfamily members in multipass membrane protein biogenesis has been observed in bacteria as well, where in some organisms YidC is recruited to the ribosome to cooperate with SecYEG in insertion (Sachelaru et al., 2013; Wickles et al., 2014).

These observations however raise questions as to why the biogenesis machinery has expanded from bacteria to mammals. Indeed, molecular dynamics simulations suggest that given sufficient time, even multipass membrane proteins can fold autonomously (Niesen et al., 2020). Though the increased size of the membrane proteome may contribute, it is more likely that a decrease in error tolerance in multicellular organisms is the driver for evolution of additional biogenesis components. Unlike a bacterium or yeast cell, mammals rely on post-mitotic cells, including neurons and cardiomyocytes, that must persist for the entire lifespan of the organism. In these cells, the risk of cytotoxicity from misfolding or aggregation of nascent proteins required evolution of more stringent mechanisms to protect the proteostasis of the cell and thereby the health of the organism. As a result, the insertion, folding and assembly of membrane proteins must occur with even greater efficiency to avoid premature clearance by quality control machinery. These additional insertases fulfill this role, by increasing the efficiency of membrane protein biogenesis such that it can occur even in the face of robust competing degradation pathways.

## ACKNOWLEDGEMENTS

We thank Songye Chen and Oliver Clarke for technical assistance, T. Stevens and M. Hazu and all members of the Voorhees lab for thoughtful discussion. We thank the Caltech Flow Cytometry facility, the Caltech Cryo-EM facility, and the Caltech Millard and Muriel Jacobs Genetics and Genomics Laboratory. We thank Ville Paavilainen for Apratoxin A. Cryo-electron microscopy was performed in the Beckman Institute Center for TEM at Caltech, and data was processed using the Caltech High Performance Cluster, supported by a grant from the Gordon and Betty Moore Foundation. This work was supported by: the Heritage Medical Research Institute (RMV), the NIH’s National Institute Of General Medical Sciences DP2GM137412 (RMV), the Burrough’s Wellcome Fund (RMV), an NSF-CAREER award 2145029 (RMV), a Human Frontier Science Program Fellowship 2019L/LT000858 (AG), the Deutsche Forschungsgemeinschaft (TP), the Tianqiao and Chrissy Chen Institute (TP), a Rosen Family fellowship (KRP), and an Arie Jan Haagen-Smit Fellowship (KRP).

## Materials and Methods

### Plasmids and antibodies

The sequences used in cell-based assays and structural analysis were derived from UniProtKB/Swiss-Prot. These include: SEC61β (SEC61B; NP_006799.1), squalene synthase isoform 1 (SQS/FDFT1; Q6IAX1), vesicle associated membrane protein 2 (VAMP2; P63027-1), type-2 angiotensin II receptor (AGTR2; P50052), SARS-CoV-2 Membrane protein (VME1_SARS2; P0DTC5), SARS-CoV-2 ORF3a (AP3A_SARS2; P0DTC3), kappa-type opioid receptor (OPRK1; P41145), alpha-1A adrenergic receptor (ADRA1A; P35348), translocating chain-associated membrane protein 2 (TRAM2; Q15035), excitatory amino acid transporter 1 (SLC1A3/EAAT1; P43003), guided entry of tail-anchored proteins factor CAMLG (GET2/CAMLG; P49069), Yip1 domain family member 1 (YIPF1; Q9Y548), Asialoglycoprotein receptor 1 (ASGR1; P07306), trace amine-associated receptor 5 (TAAR5; O14804); melatonin receptor type 1A (MTR1A/ MTNR1A; P48039), beta-1 adrenergic receptor (ADRB1; P08588), cholecystokinin receptor type A (CCKAR; P32238), adrenocorticotropic hormone receptor (MC2R/ACTHR; Q01718), melatonin receptor type 1B (MTNR1B/MTR1B; P49286), apelin receptor (APLNR/APJ; P35414), muscarinic acetylcholine receptor M1 (CHRM1/ACM1; P11229), G-protein coupled receptor 15 (GPR15; P49685), tachykinin receptor 1 (TACR1/NK1R;), translocon-associated protein subunit alpha (SSR1/SSRA/TRAPA; P43307), ER membrane protein complex subunit 7 (EMC7; Q9NPA0), ER membrane protein complex subunit 5 (EMC5/MMGT1; Q8N4V1), ER membrane protein complex subunit 2 (EMC2; Q15006), Nicalin (NCLN; Q969V3), transmembrane protein 147 (TMEM147; Q9BVK8), Nodal modulator (NOMO2; Q5JPE7), and mannosyl-oligosaccharide 1,2-alpha-mannosidase IA (MAN1A1; P33908).

The 2nd generation lenti-viral packaging plasmid psPAX2 (Addgene plasmid #12260) was a gift from Didier Trono. The 2nd generation lenti-viral packaging plasmid pCMV-VSV-G was a gift from Bob Weinberg (Addgene plasmid # 8454). The pHAGE2 lenti-viral transfer plasmid was a gift from Magnus A. Hoffmann and Pamela Bjorkman. For inducible expression in K562 cells during CRISPRi screens, the SFFV-tet3G backbone was used (Jost et al., 2017). Though mCherry and EGFP variants were used throughout the study, they are referred to as RFP and GFP, respectively, for clarity. The GFP:RFP reporter system for reporter assays was used as previously described to assess substrate insertion (Guna et al., 2022; Inglis et al., 2020).

For expression in K562 cells during genome-wide CRISPRi screens, AGTR2 and SARS-CoV-2 ORF3a were cloned as N-terminal fusions to GFP, followed by a viral 2A skipping sequence, and RFP. For SARS-CoV-2 M, the reporter was designed using the split GFP system (Cabantous et al., 2005; Kamiyama et al., 2016). Here, the GFP11 tag (RDHMVLHEYVNAAGIT) was inserted at the N-terminal separated by a 3X-GS linker to allow for complementation with GFP1-10. The M protein was designed as a split GFP reporter while AGTR2 and ORF3a were designed to contain full GFP fusions. The latter two substrates are unstable in cells with the additional length of GFP11 fused to the N-termini. Additionally, we note that in the arrayed screen in Figure 2, all substrates except for SEC61β contain full GFP or RFP fusions. The SEC61β reporter has been previously described (Guna, Page, et al., 2023). Briefly, the TMD and flanking regions were inserted downstream of the first 70 residues of the flexible cytosolic domain of SEC61β. At the C-terminal of Sec61β, the GFP11 tag (RDHMVLHEYVNAAGIT) was inserted, separated by a 2X-GS linker. To express GFP1-10 in the ER lumen, the human calreticulin signal sequence was appended to the N-terminal of GFP1-10-KDEL as previously described (Guna et al., 2022; Pleiner et al., 2020).

Programmed dual sgRNA guide plasmids were used in assays involving depletion of two genes (Replogle et al., 2020). The following sgRNA protospacer sequences were used to generate dual guide plasmids:

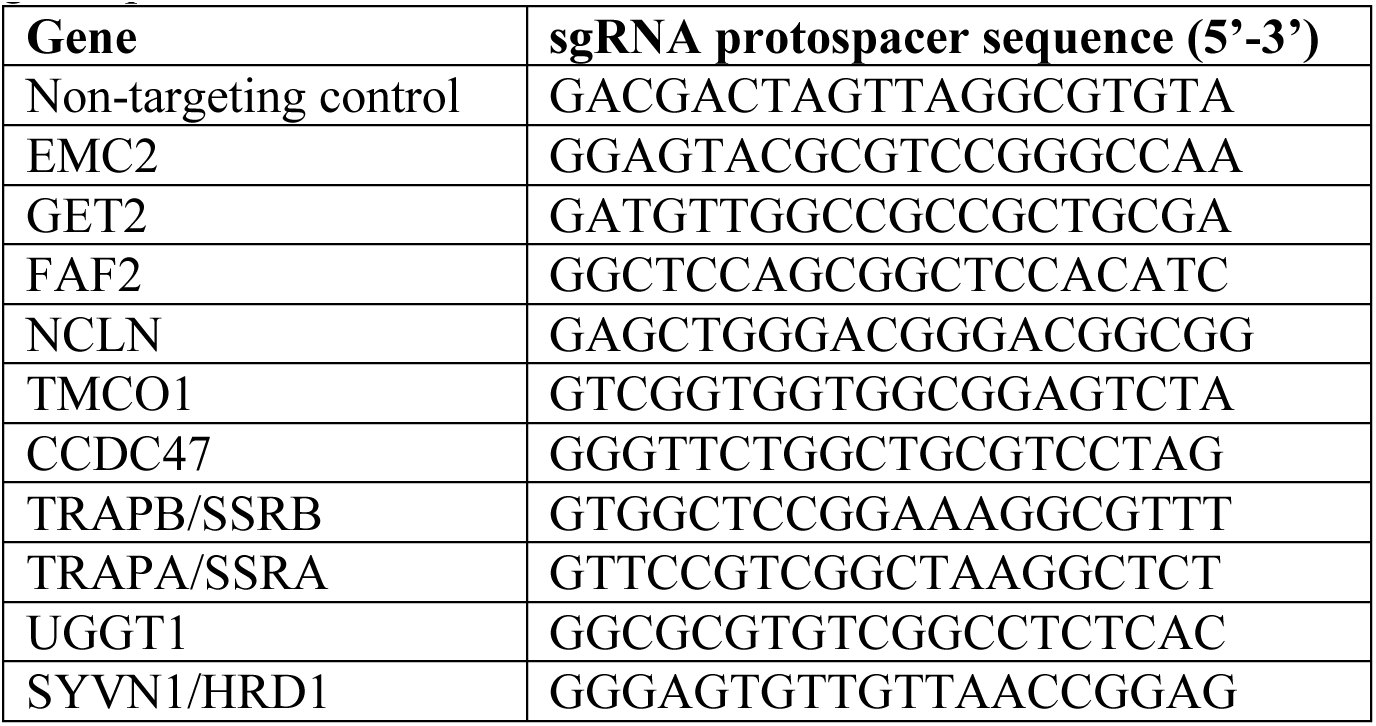

To generate knockout cell lines, the following sgRNAs were cloned into pX459 following a standard protocol: TMCO1 (GAAACAATAACAGAGTCAGC), CCDC47 (GTCGCATTCTACGTAACGGA), TMEM147 (ATGTCCCGGAATGCCGGCAA).

The following siRNAs were used in this study: negative control no. 2 siRNA (#4390846), EMC5 siRNA s41131, NOMO siRNA s49380, CCDC47 siRNA s32577 (all Silencer Select; ThermoFisher Scientific, USA).

Constructs for expression in rabbit reticulocyte lysate (RRL) were based on the SP64 vector (Promega). For *in vitro* translations, the ALFA epitope (PSRLEEELRRRLTEP) was appended to the C-terminal of SARS-CoV-2 ORF3a and M proteins,, separated by a flexible 3X-GS linker (Götzke et al., 2019).

All plasmids are available upon request.

The following antibodies were used in this study: EMC2 (25443-1-AP, Proteintech, USA); EMC3 (67205-1-Ig, Proteintech, USA); EMC4 (27708-1-AP, Proteintech, USA); EMC5 (A305-833, Bethyl Laboratories, USA); EMC7 (27550-1-AP, Proteintech, USA); EMC10 (ab180148, Abcam, UK); TMCO1 (27757-1-AP, Proteintech, USA); TMEM147 (PA5-95876, Invitrogen, USA); NCLN (A305-623A-M, Bethyl, USA); NOMO1 (PA5-47534, Invitrogen, USA); NOMO2 (PA5-95819, Invitrogen, USA); CCDC47 (A305-100A-M, Bethyl, USA); GET2 (Synaptic Systems, Germany); SYVN1 (13473-1-AP, Proteintech, USA); FAF2 (16251-1-AP, Proteintech, USA); UGGT1 (sc-374565, Santa Cruz Biotechnology, USA); Asterix (HPA06568S, Sigma Aldrich); Calnexin (10427-2-AP, Proteintech); BIP (#610979, BD Biosciences); PDI (ADI-SPA-891-D, Enzo, USA); alpha tubulin (T9026, Millipore Sigma, USA); Anti-HA-HRP (Millipore-Sigma, USA); Anti-FLAG-HRP (Millipore-Sigma, USA). The rabbit polyclonal antibodies against BAG6, TRAPA, and GFP were gifts from Ramanujan Hegde. Secondary antibodies used for Western blotting were goat anti-mouse- and anti-rabbit-HRP (#172-1011 and #170-6515, Bio-Rad, USA).

### Cell culture and cell line construction

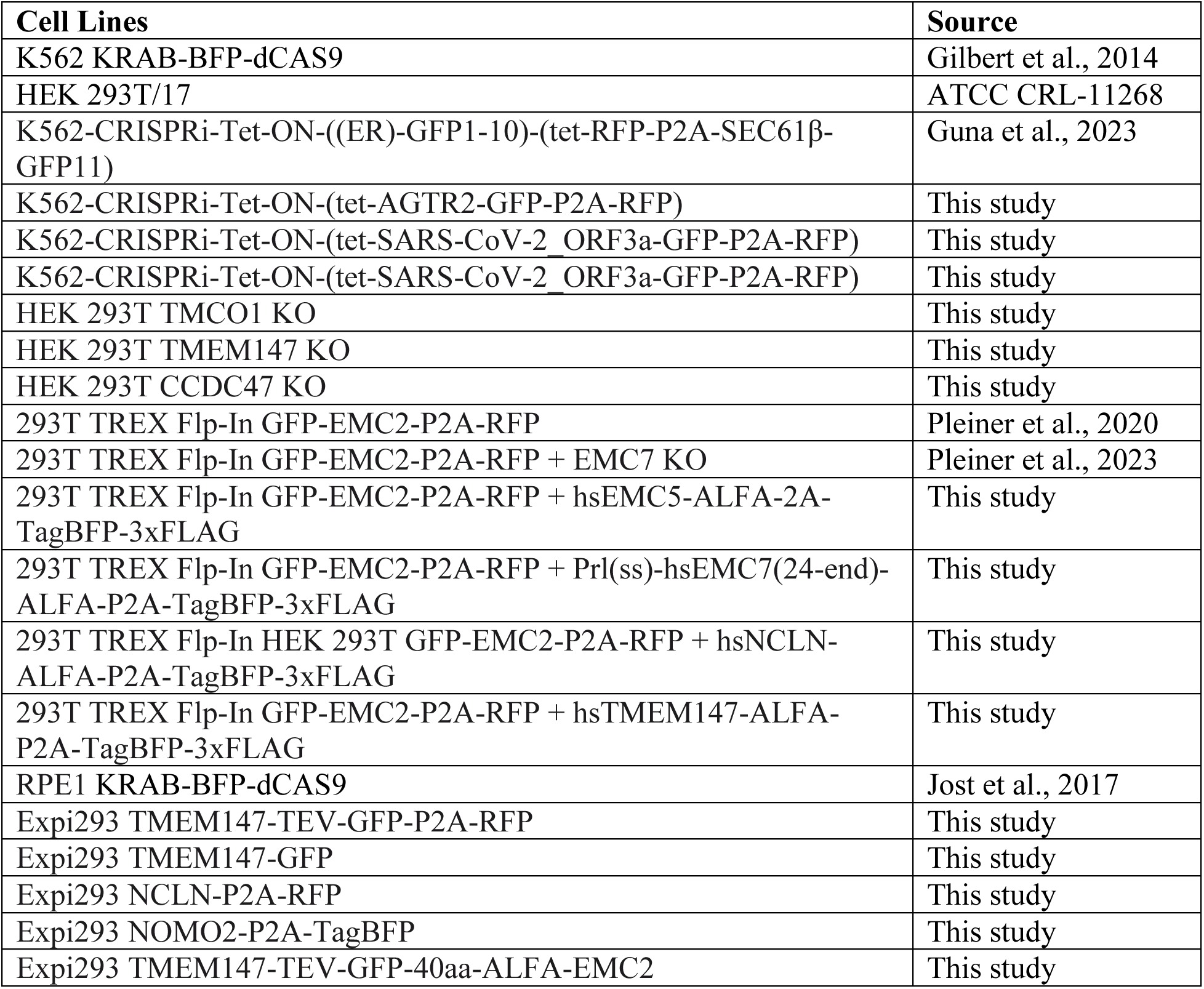

| **Cell Lines** | **Source** |
| --- | --- |
| K562 KRAB-BFP-dCAS9 | Gilbert et al., 2014 |
| HEK 293T/17 | ATCC CRL-11268 |
| K562-CRISPRi-Tet-ON-((ER)-GFP1-10)-(tet-RFP-P2A-SEC61β-GFP11) | Guna et al., 2023 |
| K562-CRISPRi-Tet-ON-(tet-AGTR2-GFP-P2A-RFP) | This study |
| K562-CRISPRi-Tet-ON-(tet-SARS-CoV-2 ORF3a-GFP-P2A-RFP) | This study |
| K562-CRISPRi-Tet-ON-(tet-SARS-CoV-2 ORF3a-GFP-P2A-RFP) | This study |
| HEK 293T TMCO1 KO | This study |
| HEK 293T TMEM147 KO | This study |
| HEK 293T CCDC47 KO | This study |
| 293T TREX Flp-In GFP-EMC2-P2A-RFP | Pleiner et al., 2020 |
| 293T TREX Flp-In GFP-EMC2-P2A-RFP + EMC7 KO | Pleiner et al., 2023 |
| 293T TREX Flp-In GFP-EMC2-P2A-RFP + hsEMC5-ALFA-2A-TagBFP-3xFLAG | This study |
| 293T TREX Flp-In GFP-EMC2-P2A-RFP + Prl(ss)-hsEMC7(24-end)-ALFA-P2A-TagBFP-3xFLAG | This study |
| 293T TREX Flp-In HEK 293T GFP-EMC2-P2A-RFP + hsNCLN-ALFA-P2A-TagBFP-3xFLAG | This study |
| 293T TREX Flp-In GFP-EMC2-P2A-RFP + hsTMEM147-ALFA-P2A-TagBFP-3xFLAG | This study |
| RPE1 KRAB-BFP-dCAS9 | Jost et al., 2017 |
| Expi293 TMEM147-TEV-GFP-P2A-RFP | This study |
| Expi293 TMEM147-GFP | This study |
| Expi293 NCLN-P2A-RFP | This study |
| Expi293 NOMO2-P2A-TagBFP | This study |
| Expi293 TMEM147-TEV-GFP-40aa-ALFA-EMC2 | This study |

K562 cells containing KRAB-BFP-dCas9 (Gilbert et al., 2014) were cultured in RPMI-1640 with 25 mM HEPES, 2.0 g/L NaHCO3, and 0.3 g/L L-glutamine supplemented with 10% Tet System Approved FBS, 2 mM glutamine, 100 units/mL penicillin, and 100 μg/mL streptomycin. K562 cells were maintained between 0.25 × 10^6^ –1 × 10^6^ cells/mL. HEK 293T cells were cultured in Dulbecco’s Modified Eagle Medium (DMEM) supplemented with 100 units/mL penicillin, 100 μg/mL streptomycin, and 10% FBS. RPE1 cells containing the KRAB-BFP-dCas9 machinery (Jost et al., 2017) were cultured in DMEM F12 medium supplemented with 10% FBS and 2 mM glutamine. K562, HEK 293T, and RPE1 cells were grown at 37 °C. Expi293 cells were cultured in Expi293 Expression Medium (Gibco) supplemented with 10% FBS and 2 mM glutamine. Expi293 cells were maintained between 0.5 × 10^6^ – 2 × 10^6^ cells/mL and harvested at 6 × 10^6^ cells/mL.

Clonal knockouts of TMCO1, CCDC47 and TMEM147 were obtained by transfecting HEK 293T cells with pX459 encoding the respective sgRNA using TransIT-293 transfection reagent (Mirus, USA). 72 h post transfection, single cells were sorted into 96-well plates using a SONY cell sorter (SH800S), and clones were selected following verification of protein depletion by Western blotting.

### Fluorescent reporter CRISPRi screens

CRISPRi screens were performed as previously described, with minor modifications (Gilbert et al., 2014; Horlbeck et al., 2016). For AGTR2 and Sec61β, screens were performed using either the Non-targeting-dual guide library or the EMC2-dual library. For SARS-CoV-2 M and ORF3a, screens were performed with the single CRISPRi-v2 library. We have previously demonstrated that the additional non-targeting guide in the dual guide cassette does not appreciably alter knockdown efficiency of the second guide in the cassette (Guna, Page, et al., 2023). CRISPRi libraries (single CRISPRi-v2 library, Non-targeting dual library [Addgene #197348], or EMC2 dual library [Addgene #197349]) were transduced at a multiplicity of infection less than one into 300-330 million K562-CRISPRi-Tet-ON cells containing the appropriate reporter. For the duration of the screens, cells were maintained in 1L spinner flasks (Bellco, SKU: 1965-61010) at a volume of 1L. 48 hours post-transfection, BFP positive cells were between 30-40%. After 48h, cells were treated with 1 µg/mL puromycin for three days to select for guide positive cells. Following approximately two days of recovery after puromycin selection, the reporter was induced with doxycycline (100-1000 ng/mL) for 24-48 hours and sorted on a FACSAria Fusion Cell Sorter. Cells were diluted to 0.5 × 10^6^ cells/mL each day to ensure that the culture was maintained at an average coverage of more than 1000 per sgRNA.

For sorting, cells were gated for BFP (selecting guide-positive cells), RFP and GFP (selecting an expressing reporter). Cells were sorted based on the GFP:RFP ratio of the final gated population. Approximately 30 million cells with either the highest or lowest 30% GFP:RFP ratios were collected during sorting, pelleted, and flash-frozen. From cell pellets, genomic DNA was extracted and purified using a Nucleospin Blood XL kit (Takara Bio, #740950.10). The guides were amplified and barcoded by PCR using NEB Next Ultra ii Q5 MM (M0544L). For both single and dual guide CRISPRi screens, a unique forward index primer was used. For single guide CRISPRi screens, a reverse primer that binds downstream of the guide was used (5’- CAAGCAGAAGACGGCATACGAGATCGACTCGGTGCCACTTTTTC). For dual guide CRISPRi screens, a reverse primer that binds in the hU6 region upstream of the fixed guide was used (5’-CAAGCAGAAGACGGCATACGAGATGGAATCATGGGAAATAGGCCCTC), as previously described (Guna et al., 2023). SPRISelect beads (Beckman Coulter B23317) were used to purify the DNA library (279 or 349 bp), and purified DNA was analyzed on an Agilent 2100 Bioanalyzer prior to sequencing using an Illumina HiSeq2500 using the standard CRISPRi-v2 library sequencing primer (5’- GTGTGTTTTGAGACTATAAGTATCCCTTGGAGAACCACCTTGTTG). Analysis of the sequencing was performed using the pipeline in https://github.com/mhorlbeck/ScreenProcessing <u>(Horlbeck et al., 2016)</u>. To ensure coverage, guides with fewer than 50 counts were excluded from analysis. The phenotype score for each gene was calculated from the strongest 3 sgRNA phenotypes. The Mann-Whitney p-value was calculated using the 5 sgRNAs targeting the same gene compared to the negative controls. For screens that were performed in biological duplicate (SARS-CoV-2 M, SARS-CoV-2 ORF3a, and Sec61β), the sgRNA phenotypes were averaged. To calculate the discriminant scores used in Fig. 3, each gene’s phenotype score was multiplied by its Mann-Whitney p-value.

### Lentiviral Transduction

Lentivirus was generated via co-transfection of a transfer plasmid of interest along with packaging plasmid psPAX2 and envelope plasmid VSV.G, using TransIT-293 transfection reagent (Mirus). Lentivirus was harvested 48 hours after transfection, then aliquoted, flash-frozen, and stored for future usage.

Cell lines for structural analysis were generated in Expi293 cells. Suspension cells were transduced by mixing 10 million cells with 2.5 ml of harvested lentiviral supernatant in presence of 8 μg/ml polybrene in a final volume of 30 ml in a 125-ml vented flask. For BOS (tNOMO) cell line, 2.5 ml of lentiviral supernatant of each subunit TMEM147-GFP, NCLN-P2A-RFP, NOMO(Δ1-9Ig)-P2A-TagBFP were added during transduction. The cells were grown in a shaking incubator for ∼16 hours before being pelleted and resuspended in 50 ml of fresh medium in order to remove lentiviral particles. Then the cells were continued to be grown for about a week until transduced cells expressing plasmid of interest were sorted with the Sony SH800S cell sorter (Sony Biotechnology).

K562 dCas9-BFP-KRAB cells were spinfected with lentivirus containing dual sgRNAs targeting two genes of interest or a non-targeting control. Briefly, 250,000 cells were mixed with 200 µl of lentivirus and RPMI medium in the presence of 8 µg/ml polybrene in a total volume of 2 ml in a 12-well plate. Plates containing K562 cells were spun at 1,000 g for 1.5-2 h at 30°C, resuspended, and cultured in 12-well plates. Approximately 48 h after spinfection, 1 µg/ml puromycin was added for 5 consecutive days to select cells containing the dual guide cassette. To assess the percentage of guide-containing, BFP-positive cells, samples were analyzed using flow cytometry, as described below. After a total of 8 days of knockdown, cells were pelleted, flash frozen, and used in western blot analysis to assess knockdown of individual genes.

For reporter assays in adherent HEK293 or RPE1 cells, lenti-viral transduction of 50-300 µl lentiviral supernatant and 8 µg/ml polybrene (Millipore-Sigma, USA) were added to ∽70% confluent cells in 2.5 ml culture medium in a 6-well. Lenti-viral reporter constructs of all GPCRs, TRAM2, SARS-CoV-2 M, SARS-CoV-2 ORF3a, EAAT1, GET2, YIPF1 for use in HEK 293T and RPE1 cells, contained an upstream CMV promoter followed by the protein fused to GFP, a 2A site, and RFP (Figures 2B,C, 3D-F, 4C, 6A, 6C-F, S6A, C-F, S3C-D, S7A). Versions of GPCRs with a signal sequence (Figures 4C, 6C, and S6C) contained N-terminal fusions of the pre-prolactin signal sequence (KGSSQKGSRLLLLLVVSNLLLCQGVV) followed by a T4 Lysozyme soluble domain. In parallel, the first TMD (residues 33-75) of MAN1A1, a membrane protein with N_cyt_ topology, was fused to the N-terminus of GPCRs. Both signal sequence-T4 lysozyme fusions and MAN1A1 fusions behaved similarly (Figure 4C and S3D). Sec61β, SQS, VAMP2, and ASGR1 lenti-viral reporters for use in HEK 293T and RPE1 cells contained an upstream CMV promoter, followed by GFP, a 2A site and RFP, which was fused to the reporter. The TMD and flanking regions of Sec61β, SQS, and VAMP2 the were fused directly to RFP, as described before (Guna et al., 2018; Guna, Page, et al., 2023; Pleiner et al., 2020). A charge mutant of OPRK1 (E45K, D46R, E50K) (+0 variant) was used in RPE1 cells in the same GFP-2A-RFP cassette as described above for GPCRs and as previously described (Pleiner et al., 2023) (Figures 6C-F, S6C, S7A).

For CRISPRi knockdown experiments in RPE1 cells, cells were transduced with sgRNA dual guide lenti-viral vectors. After 6 days of knockdown, cells were transduced with fluorescent reporter lenti-viral vectors described above and analyzed ∼48h post-transduction (8 days after transduction with guide).

For rescue assay experiments, 300,000 HEK 293T and HEK 293T EMC7 KO cells were seeded into each 6-well plate on Day 1. On Day 2, cells were transduced with 300 μl lentiviral supernatant of rescue construct(s) and 8 μg/ml final concentration of polybrene, marking the start of the 72-hour rescue lentivirus addition. The media was exchanged on Day 3 to remove excess polybrene, and the 48-hour reporter lentivirus addition started by transducing the cells with 150 μl lentiviral supernatant of reporter construct(s) in presence of 8 μg/ml final concentration of polybrene. On Day 4, the cells were split 1:2 into a different set of 6-well plates to be used for Western Blot. Lastly, on Day 5, the cells were harvested, washed and resuspended in 500 μl Dulbecco’s Phosphate Buffered Saline (Gibco) to be analyzed by flow cytometry or frozen for analysis via Western Blot.

### Flow cytometry

RPE1 and HEK 293T cells were trypsinized, washed with 1xPBS, and resuspended in 1xPBS for flow cytometry analysis. K562 cells were analyzed directly from 12-well or 6-well cultures. Cells were analyzed using an Attune NxT Flow Cytometer (Thermo Fisher Scientific, USA) or a MACSQuant VYB (Miltenyi Biotec, Germany). Flow cytometry data was analyzed using FlowJo v10.8 Software (BD Life Sciences, USA) or by Python using the FlowCytometryTools package.

The Sec61 inhibitor Apratoxin A was used to analyze the effect of SEC61 inhibition on membrane protein insertion (Paatero et al., 2016; Thornburg et al., 2013). HEK 293T (WT or TMCO1 KO) cells were transduced with reporter lenti-virus, and 48h later, cells were treated with 31.3 nM Apratoxin A in 0.1% DMSO for 12h. Cells were analyzed immediately following treatment with inhibitor. Apratoxin A was a gift from Ville Paavilainen.

### Preparation of human ER microsomes

Human derived rough ER microsomes were generated as previously described, with minor modifications (Chitwood et al., 2018). HEK293T cells (WT, NCLN KO, TMCO1 KO, or EMC6 KO) were harvested and washed in 1X PBS. Cells were resuspended in 4 times the pellet volume of sucrose buffer (10 mM HEPES, pH 7.5, 250 mM sucrose, 2 mM magnesium acetate, 1X cOmplete EDTA-free protease inhibitor cocktail [Roche]) and lysed by douncing at 4 °C. Lysed cells were diluted 2X in sucrose buffer and pelleted at 3214 xg for 35 min. at 4 °C. Supernatant was transferred to a new tube and pelleted again at 3214 xg for 35 min. at 4 °C. To isolate the microsomal fraction, samples were pelleted in an ultracentrifuge in an MLA80 rotor (Beckman-Coulter) at 75,000 xg for 1h at 4 °C. Supernatant was removed, and the microsomal pellet was resuspended to an A280 of 75 in microsome buffer (10 mM HEPES, pH 7.5, 250 mM sucrose, 1 mM magnesium acetate, 0.5 mM DTT). To remove contaminating RNAs, microsomes (hRMs) were nucleased. CaCl_2_ (1 mM) and micrococcal nuclease (0.125 U/μL) were added to hRMs and mixed before incubating for 6 minutes at 25 °C. To quench the reaction, EGTA (2 mM) was added to the sample and the sample was immediately mixed and place on ice. Nucleased hRMs were flash frozen and stored at -80 °C prior to use in *in vitro* translations.

### Mammalian in vitro translation

Translation extracts were prepared using nucleased rabbit reticulocyte lysate (RRL) supplemented with human derived rough ER microsomes, as previously described (Sharma et al., 2010; Walter & Blobel, 1983). DNA templates for *in vitro* transcription were made by PCR from SP64-based plasmids or directly from double-stranded DNA gene fragments (IDT or Twist Biosciences) using primers within the SP6 promoter (5’ end) and following a stop codon and short untranslated region (3’ end). Run-off transcription reactions were made by combining 4.8 μL T1 mix (Sharma et al., 2010), 0.1 µL RNasin (Promega), 0.1 µL SP6 polymerase (New England Biolabs) and 50 ng PCR product. Reactions were incubated at 37 °C for 2 hours, and then used directly in translation reactions, which were incubated for 20-45 minutes at 32 °C. To label nascent proteins, radioactive ^35^S-methionine (Perkin Elmer) was included in translation reactions, unless otherwise indicated. Samples were then analyzed directly using SDS-PAGE and autoradiography.

For experiments in which the insertion of the first TMD was assessed, substrates were translated in the presence of hRMs derived from HEK 293T cells. The OPRK1 constructs (wildtype or variants with 3xK or 5xK mutations in the N-terminal soluble domain) contain an Asn-Gly-Thr (NGT) glycosylation site at the N-terminus, which allows monitoring of insertion. The VAMP2 control protein contains a C-terminal Opsin tag that gets glycosylated upon insertion of the TA substrate and allows monitoring of insertion. A construct containing the first 85 amino acids of preprolactin was used as a control for signal sequence cleavage. For assays in which Sec61’s insertion capacity was assessed, the inhibitor Apratoxin A was used at 1 μM.

### Preparation of the ALFA nanobody conjugated to HRP for Western blotting

The ALFA nanobody was coupled to HRP-maleimide through a single engineered C-terminal cysteine residue, as previously described (Pleiner et al., 2018).

### DSP crosslinking

Suspension adapted T-REx-293 cells stably expressing either GFP-EMC2 only or GFP-EMC2 plus EMC5-ALFA, EMC7-ALFA, TMEM147-ALFA, or NCLN-ALFA were harvested, washed in PBS, pelleted, and resuspended in PBS containing 1.5 mM final concentration of dithiobis(succinimidyl propionate) (DSP; Thermo Scientific). The cell mixture was incubated at 4°C with head-over-tail rotation for 2 hours. After the incubation, the reaction was quenched by addition of 1M Tris/HCl, pH 7.5 to 20 mM final concentration and incubated for 15 min. Then, the cells were pelleted, weighed, and flash frozen for storage prior to immunoprecipitation or prepared for mass spectrometry, as described below.

LC-MS/MS analysis for the IP experiment was performed with an EASY-nLC 1200 (ThermoFisher Scientific, San Jose, CA) coupled to a Q Exactive HF hybrid quadrupole-Orbitrap mass spectrometer (ThermoFisher Scientific, San Jose, CA). Peptides were separated on an Aurora UHPLC Column (25 cm × 75 μm, 1.6 μm C18, AUR2-25075C18A, Ion Opticks) with a flow rate of 0.35 μL/min for a total duration of 75 min and ionized at 1.6 kV in the positive ion mode. The gradient was composed of 6% solvent B (3.5 min), 6-25% B (42 min), 25-40% B (14.5 min), and 40–98% B (15 min); solvent A: 2% ACN and 0.2% formic acid in water; solvent B: 80% ACN and 0.2% formic acid. MS1 scans were acquired at the resolution of 60,000 from 375 to 1500 m/z, AGC target 3e6, and maximum injection time 15 ms. The 12 most abundant ions in MS2 scans were acquired at a resolution of 30,000, AGC target 1e5, maximum injection time 60 ms, and normalized collision energy of 28. Dynamic exclusion was set to 30 s and ions with charge +1, +7, +8 and >+8 were excluded. The temperature of ion transfer tube was 275°C and the S-lens RF level was set to 60. MS2 fragmentation spectra were searched with Proteome Discoverer SEQUEST (version 2.5, Thermo Scientific) against *in silico* tryptic digested the UniProt Human proteome Swiss-Prot database (UP000005640). The maximum missed cleavages were set to 2. Dynamic modifications were set to oxidation on methionine (M, +15.995 Da), deamidation on asparagine and glutamine (N and Q, +0.984 Da) and protein N-terminal acetylation (+42.011 Da). Carbamidomethylation on cysteine residues (C, +57.021 Da) was set as a fixed modification. The maximum parental mass error was set to 10 ppm, and the MS2 mass tolerance was set to 0.03 Da. Intensity-based quantification (iBAQ) was performed using the IMP-apQuant PD node (Doblmann et al., 2019; Schwanhäusser et al., 2013). The maximum false peptide discovery rate was specified as 0.01 using the Percolator Node validated by q-value. Data of the abundance rank in each replicate are available in Table S4.

### Protein purification for structure determination

2 L of Expi293 cells stably expressing the protein(s) of interest by lentiviral transduction were pelleted, washed with PBS and flash-frozen for storage. For BOS (fNOMO), a cell line was generated stably expressing TMEM147-GFP-2A-RFP. For BOS (tNOMO), a cell line stably expressing TMEM147-TEV-GFP, NCLN-RFP, NOMO(Δ1-9Ig)-BFP. For the BOS (fNOMO) • EMC holocomplex, we generated a cell line stably expressing TMEM147-5aa-TEV-GFP-40aa(ALFA)-EMC2. All protein complexes were purified using an anti-GFP nanobody as described previously (Pleiner et al., 2020; Stevens et al., 2023). Briefly, cell pellets were harvested, washed with 1xPBS, and resuspended in solubilization buffer (50 mM HEPES/KOH pH 7.5, 200 mM NaCl, 2 mM MgAc_2_, 1x cOmplete^TM^ EDTA-free Protease Inhibitor Cocktail [Roche], 1% [w/v] glyco-diosgenin [GDN; Anatrace], 1 mM DTT) at a ratio 6.8 ml solubilization buffer per 1 g cell pellet. Following incubation for 1 hour at 4°C, lysate supernatant was isolated by centrifugation at 18,000 rpm using an SS-34 rotor in a Sorvall RC6+ Superspeed Centrifuge at 4°C for 45 min.

Simultaneously, biotinylated anti-GFP nanobody was immobilized onto streptavidin magnetic beads. Specifically, 80 μl resuspended Pierce™ Streptavidin magnetic beads per 1 g cell pellet were washed and equilibrated in wash buffer (50 mM HEPES pH 7.5, 200 mM NaCl, 2 mM MgAc_2_, 0.0053% GDN, 1 mM DTT). Then His14-Avi-SUMO^Eu1^-tagged anti-GFP nanobody (Addgene #149336) was immobilized onto the washed magnetic beads for 30 min with mixing at 4°C using a ratio of 27 μg for every 80 μl beads. This immobilization was followed by incubation of beads with 50 mM HEPES/KOH pH 7.5 containing 100 μM biotin for 5 min on ice to block unbound biotin binding sites on the magnetic streptavidin beads.

Subsequently, the beads were washed with solubilization buffer and incubated with clarified cell lysate for 1 hour with head-over-tail rotation. After incubation, 4 washes with wash buffer (2 volumes of wash buffer:1 volume of beads) was performed to remove unspecific binding to the beads. To elute the bound proteins, wash buffer containing 500 nM SUMO^Eu1^ protease (Addgene #149333) was added to the beads and left to incubate for 30 minutes with mixing at 4°C. The eluent was further purified using size exclusion chromatography with a 3.5 ml Superose 6 column (GE Life Sciences). For BOS (tNOMO) sample, TEV protease was added (1 mg TEV protease for every 30 mg of BOS (tNOMO) protein) and incubated overnight at 4°C without mixing to remove the GFP-tag before size exclusion chromatography. The fractions corresponding to the protein complexes were concentrated using a 500-μl 30K MWCO concentrator (Millipore-Sigma).

### Grid preparation and data collection

For BOS (fNOMO) sample, 3 μL of purified, concentrated protein at 2.48 mg/ml was applied to UltrAuFoil® R 1.2/1.3 holey gold film grid (Ted Pella, Inc.) that had been grow discharged with the PELCO easiGlow^TM^ (Ted Pella, Inc.) at 20 mA for 60 s. The grid was blotted at 6°C, 100% humidity, -4 blot force for 4 seconds and plunged frozen in liquid ethane using the FEI Vitrobot Mark IV (Thermo Fisher Scientific). Data were collected on a Titan Krios operating at 300 keV and equipped with a Gatan K3 direct detector and a 20 eV slit width energy filter. Images were acquired using an automated acquisition pipeline in SerialEM (Mastronarde, 2005) and recorded at 105k magnification with a defocus range of -3.0 to -1.0 μm and total exposure dose of 60 e^-^/Å^2^ in super resolution mode with a pixel size of 0.418 Å/pixel. 11,870 micrographs were collected for this data set.

For BOS (tNOMO), the grid was prepared in a similar manner, except the protein sample was concentrated to 4 mg/ml and mixed with 0.005% 3-([3-Cholamidopropyl]dimethylammonio)-2-shydroxy-1-propanesulfonate (CHAPSO; Sigma Aldrich) immediately before vitrification. 15,929 micrographs were collected for this data set.

For the BOS (fNOMO) • EMC homocomplex sample, the protein concentration was at 2.53 mg/ml and 17,978 micrographs were collected for this data set.

### Structure image processing

The workflows for data processing of BOS (fNOMO), BOS (tNOMO), and BOS (fNOMO) • EMC are summarized in Fig. S5 A, B, C, respectively. Data processing was carried out using cryoSPARC v3.2-4.2.1 (Punjani et al., 2017). For preprocessing, micrographs were motion-corrected, Fourier-cropped twofold to 0.832 Å/pixel using ‘Patch Motion Correction’; then, they were subjected to patch-based contrast transfer function (CTF) estimation with ‘Patch CTF Estimation’. Movies were selected based on CTF fit cut-off of 5.0 Å in ‘Curate Exposure’. From here on out, details of data processing differ for each structure.

For BOS (fNOMO), 814,566 particles were picked using ‘Blob Picker’ and extracted with box size = 512 pixels from 7,174 selected movies. Iterative rounds of ‘2D classifications’ performed to remove background and junk particles. 2D classes that resemble BOS (fNOMO) complex were used as template for ‘Template Picker’ with particle diameter = 190 Å, which resulted in 904,456 picked particles. A round of ‘2D Classification’ was performed to remove background particles, resulting in 246,296 particles, which were subjected to 2 more rounds of 2D classification. Then, the resulting 90,893 particles were used to generate 4 3D classes with ‘Ab-Initio Reconstruction’. Using these 4 volumes, we performed 2 rounds of 3D ‘Heterogeneous Refinement’ on the 246,296 particles from earlier to arrive at the final EM map of BOS (fNOMO), generated from a set of 63,018 particles.

For BOS (tNOMO), 1,900,000 particles were picked using ‘Blob Picker’ with particle diameter of 120-320 Å and extracted from micrographs using box size = 512 pixels from 13,196 selected movies. After iterative rounds of ‘2D Classification’, 319,567 particles were used to generate a 3D volume using ‘Ab-Initio Reconstruction’. This volume was then used for template generation for template picking, resulting in 2,500,000 picked particles. After iterative rounds of 2D classification, 1,265,788 particles were subjected to ‘Ab-Initio Reconstruction’ into 4 volumes. The particles that correspond to the 2 volumes that best resembled our BOS (tNOMO) complex were put through multiple rounds of 3D ‘Heterogeneous Refinement’ to give us 312,033 particles which were re-extracted using box size of 448 pixels. These particles were used as input in more rounds of 3D ‘Heterogeneous Refinement’ and ‘3D Classification (BETA)’ to arrive at an EM map generated from 115,841 particles. This map was put through 3D ‘Non-uniform Refinement’ and sharpened with a B-factor of -60 Å^2^ to give us our final map. Additionally, ‘DeepEMhancer’ was also used to aid in model building.

For BOS (fNOMO) • EMC, 3,100,000 particles were picked from 17,586 selected movies using ‘Blob Picker’ with particle diameter of 175-450 Å, which were then extracted from micrographs 2x binned and subjected to 4 rounds of ‘2D Classification’ to remove background particles. 300,442 particles were used to construct 3 ab-initio models with ‘Ab-Initio Reconstruction’ (with default settings except for maximum resolution = 7, initial resolution = 9, and initial minibatch size = 300, final minibatch size = 1000). The map that best resembled EMC with BOS was used to generate 2D templates for template particle picking (‘Template Picker’) with particle diameter of 300 Å, resulting in 1,932,646 picked particles. After 1 round of ‘2D Classification’ and ‘Heterogeneous Refinement’ using 3 junk classes that resemble background and 1 class that resembles EMC with BOS, 595,637 particles were subjected to a first round of ‘Ab-initio Reconstruction’ into 2 classes (default settings except: maximum resolution = 9, initial mini batch size = 400, final mini batch size = 1200). From the 3 replicate runs of the first round of ab-intio reconstruction, the particles associated with the better 3D volume that represented BOS (fNOMO) • EMC were combined and used as input in the second round of ab-initio reconstruction into 2 classes (same settings as previous round except: maximum resolution = 6, initial resolution = 12). This process was repeated for the third time with the second rounds’ particles that correspond to the better 3D volume (similar settings except: maximum resolution = 5, initial resolution = 7). For the subsequent rounds of ab-initio reconstruction, the settings were similar, except for maximum resolution = 4, initial resolution = 6. The particles were further classified using 3D heterogeneous refinement to achieve a final class of 69,845 particles. To get our final map, local refinement was performed on this map using a mask on the BOS complex generated in Chimera (Pettersen et al., 2004) with soft padding = 12, using pose/shift gaussian prior during alignment with standard deviation of prior over rotation = 3 degree and standard deviation of prior over shifts = 1 Å.

### Model building and refinement

For the BOS (fNOMO) structure, initial models of each subunit (TMEM147, NCLN, NOMO) were generated using AlphaFold2-Multimer ColabFold (AlphaFold2_advanced.ipynb) (Mirdita et al., 2022). Since the map quality was not sufficient for accurate model building and refinement, the initial models of each subunit were only rigid body fitted into the EM density (Fig. 5 A) and combined in COOT (Casañal et al., 2020; Emsley et al., 2010).

For the BOS (tNOMO) structure, we used the previously generated models from AlphaFold2 to rigid body fit into the density. The models were combined and manually refined in COOT (Casañal et al., 2020; Emsley et al., 2010). The final model was iteratively subjected to *phenix.real_space_refinement* (Afonine et al., 2018; Liebschner et al., 2019) with rigid body and secondary structure restraints.

For BOS (fNOMO) • EMC structure, we used BOS (tNOMO) and EMC structure (PDB: 8S9S) as initial models, which were combined, rigid body fitted and refined manually in COOT (Casañal et al., 2019; Emsley et al., 2010). The model was iteratively subjected to *phenix.real_space_refinement* (Afonine et al., 2018; Liebschner et al., 2019) with rigid body and secondary structure restraints.

CryoEM data collection, refinement, and validation statistics are reported in Table S5. Final models were evaluated with MolProbity. All figures in this study were generated with PyMOL (www.pymol.org) and ChimeraX (Goddard et al., 2018; Pettersen et al., 2021).

---

**Figure S1.**
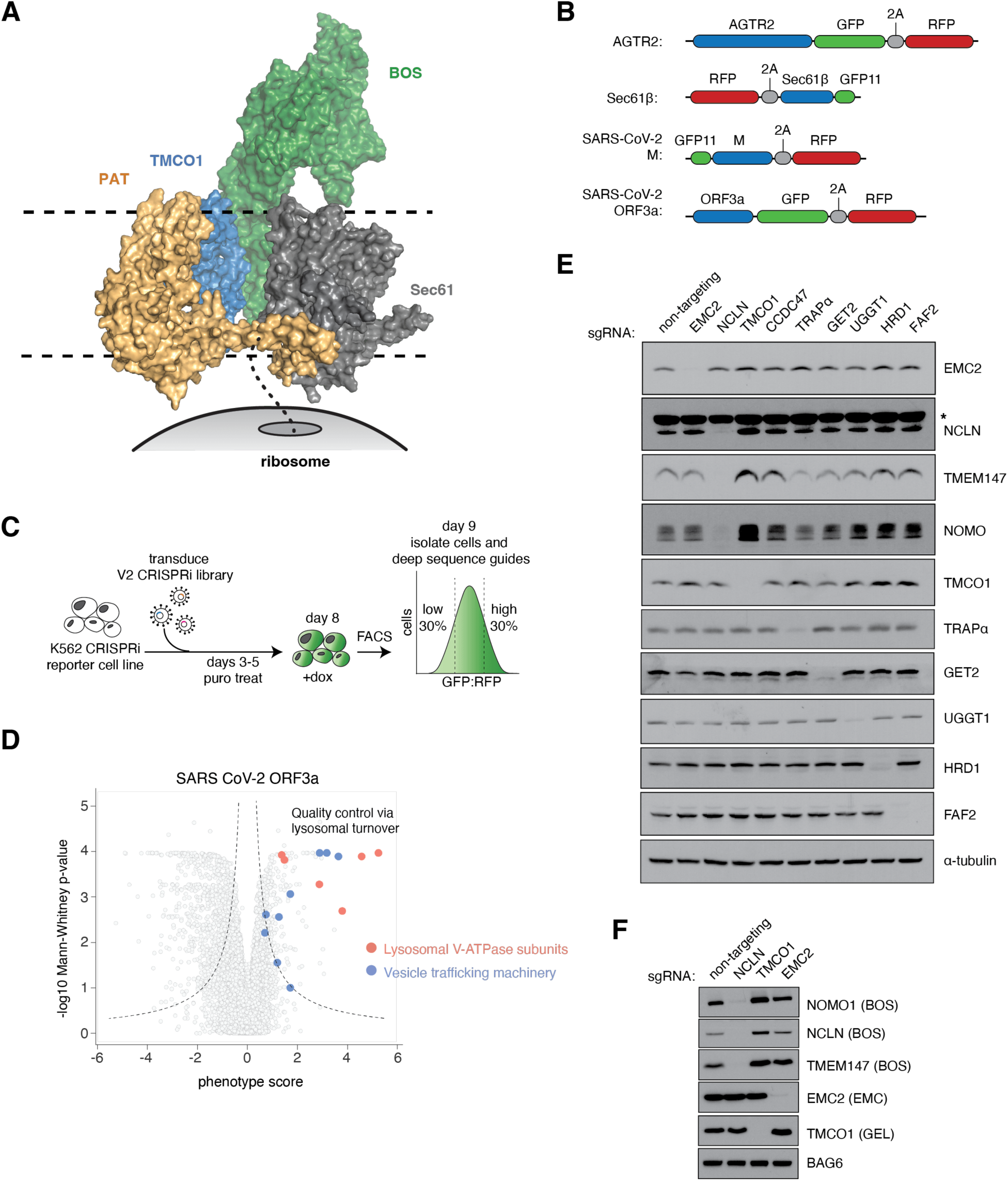
Genome-wide analysis of membrane protein biogenesis factors. **(A)** Surface representation of the ER multipass translocon (PDB ID: 6W6L) (McGilvray et al., 2020) with respect to the ribosome. SEC61 is shown in grey, BOS complex in green, TMCO1 in blue, and PAT complex in dark beige. The dotted line emerging from the ribosome represents the nascent chain of protein being inserted by the translocon. (**B**) Schematic of reporters used in CRISPRi screens. **(C)** Schematic of CRISPRi screens performed on the AGTR2, SEC61β, SARS-CoV-2 M, and SARS-CoV-2 ORF3a proteins. **(D)** Volcano plot of factors involved in the turnover of SARS-CoV-2 ORF3a protein. ORF3a is subject to degradation by the lysosome, and therefore knockdown of vesicle trafficking machinery and of the lysosomal V-ATPase subunits stabilizes the ORF3a reporter. **(E)** Analysis of gene knockdown in K562 cells by guides in CRISPRi screens. Samples were subject to SDS-PAGE and western blotting. **(F)** ER insertion machinery shows compensatory effects upon the loss of individual factors. Samples were generated and analyzed as in (E), in K562 cells. Note that loss of TMCO1 results in increased levels of NOMO, NCLN, and TMEM147, all components of the BOS complex. Additionally, loss of EMC2 results in increased levels of TMCO1, an evolutionarily related insertase within the OXA1 superfamily of insertases (Anghel et al., 2017).

**Figure S2.**
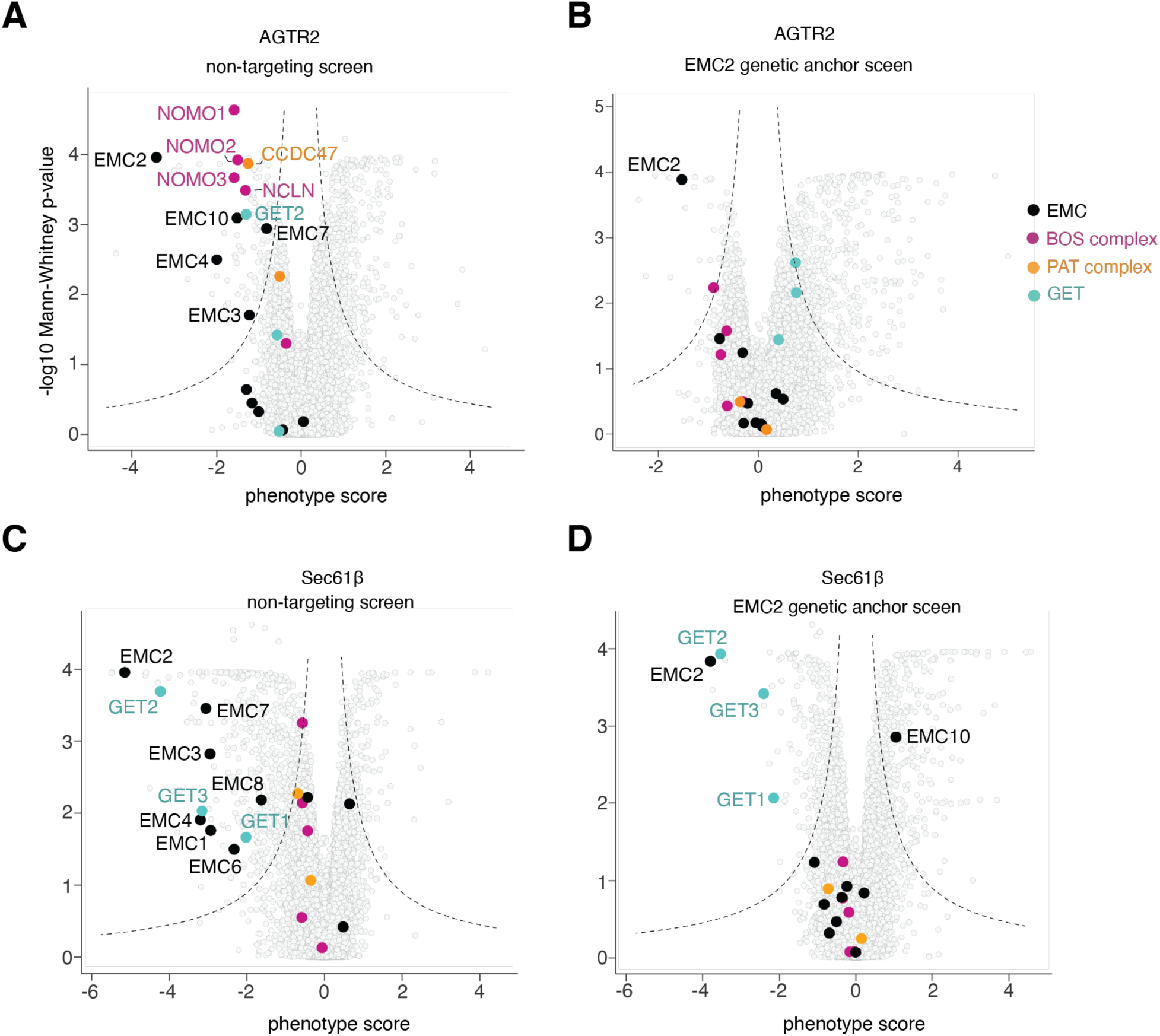
Genetic modifier screens identify epistatic and parallel pathways with the EMC. **(A)** Volcano plot of a dual guide CRISPRi screen of AGTR2 with the non-targeting (NT)-dual library. The EMC, multipass translocon components, and the GET pathway components are highlighted. Dotted lines indicate the significance of the gene’s effect on the stability of the protein reporter, and only significant gene hits from the screen are labeled. **(B)** As in (A), but with the EMC2-dual library. Note that components in the same pathway as EMC no longer have a destabilizing effect on the AGTR2 reporter. **(C)** As in (A) for Sec61β with the NT-dual library (Guna, Page, et al., 2023). **(D)** As in (A) for Sec61β with the EMC2-dual library. Note that the EMC components are no longer significant but the GET pathway components, parallel to EMC, become more significant.

**Figure S3.**
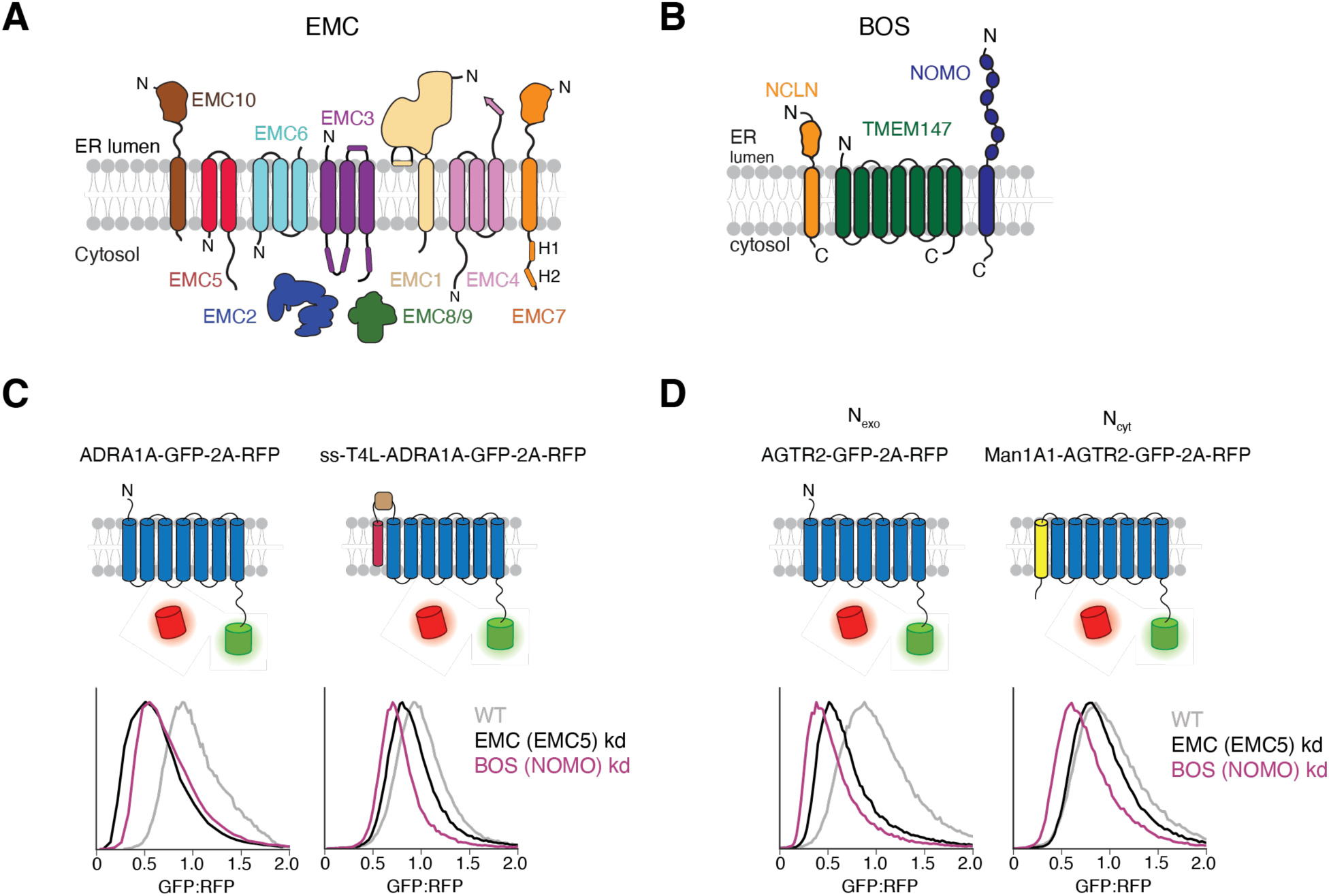
The EMC and BOS complex topologies. **(A)** Topology of the nine-subunit human EMC. In mammals, the EMC is composed of seven membrane-spanning and two soluble subunits. The central insertase is composed of the two central subunits EMC3 (a homolog of YidC and member of the Oxa1 superfamily of insertases) and EMC6 (Bai et al., 2020; Miller-Vedam et al., 2020; Pleiner et al., 2020). In both yeast and humans, the EMC also contains a globular cytosolic domain (scaffolded around EMC2 and in mammals containing the redundant paralogs EMC8 or 9) and a large lumenal domain (composed of EMC1, 7, 10, and a single β-strand of EMC4). The function of the lumenal domain of the EMC is not known, but β -propellers, like those conserved in EMC1, are typically considered protein-protein interaction motifs. **(B)** Topology of the heterotrimeric BOS complex. NOMO contains 12 lumenal IgG repeats and a single C-terminal TMD. The single-spanning protein NCLN also has a large lumenal domain, and is homologous to nicastrin, a component of the γ-secretase complex (Haffner et al., 2004). Finally, TMEM147 contains 7 TMDs and is homologous to APH-1 in the γ-secretase complex (Dettmer et al., 2010). **(C)** Flow cytometry analysis of the ratiometric ADRA1A protein reporter with or without the N-terminal fusion of the signal sequence (ss) of Pre-Prolactin followed by T4 Lysozyme (T4L), as described in Figure 4C. **(D)** Flow cytometry analysis of the ratiometric AGTR2 protein reporter with or without an N-terminal fusion of the first TMD of MAN1A1, a membrane protein of N_cyt_ topology. Note that both the N-terminal fusion of MAN1A1 and of (ss)-T4 lysozyme (Figure 4C) behave similarly and rescue the NOMO kd and EMC5 kd phenotypes.

**Figure S4.**
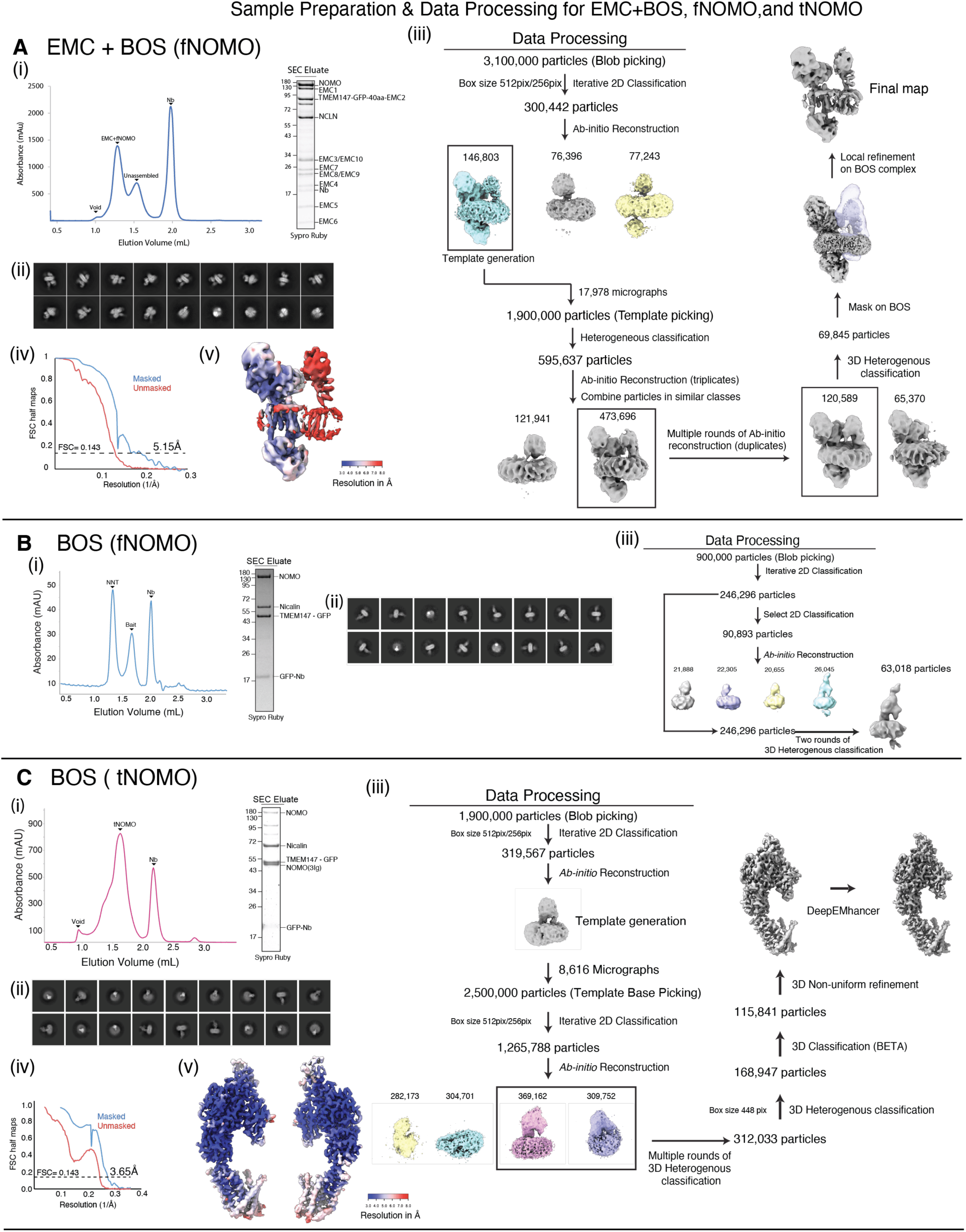
Sample preparation and data processing of EMC • BOS (fNOMO), BOS (fNOMO), and BOS (tNOMO). **(A)** EMC • BOS (fNOMO). (**i**) Sample preparation of EMC • fNOMO. Left: Size exclusion chromatogram of EMC • BOS (fNOMO). The main peak corresponds to the EMC • BOS (fNOMO) super-complex, while a smaller side peak likely corresponds to non-interacting EMC and BOS (fNOMO) complexes. Right: SDS-PAGE gel of the size exclusion eluate that contained EMC • BOS (fNOMO), stained by Sypro Ruby. The subunits of EMC and NOMO complexes are labeled. (**ii**) Representative 2D classes of EMC • BOS (fNOMO) complex. (**iii**) CryoSPARC processing scheme used to identify particles corresponding to the EMC • BOS (fNOMO) complex. CryoSPARC processing scheme employed for identifying particles related to the EMC BOS (fNOMO) complex. All processing was carried out using CryoSPARC; for details, refer to Methods (**iv**) GFSC curves for the masked and unmasked half maps of the overall EMC • BOS (fNOMO) complex map. Nominal resolution of 5.1 Å was determined using gold standard FSC cutoff = 0.143. (**v**) Resolution estimation of the EMC • BOS (fNOMO) complex, generated from cryoSPARC. **(B)** BOS (fNOMO). (**i**) Sample preparation of BOS (fNOMO). Left: Size exclusion chromatogram of BOS (fNOMO) eluant after GFP-immunoprecipitation. A280 of the run was monitored to identify peak fractions that contained BOS (fNOMO). Right: shows the SDS-PAGE gel of the purified BOS (fNOMO) complex, stained by Sypro Ruby. All the components of the BOS (fNOMO) complex were present (NOMO, NCLN, and TMEM147-GFP), along with excess anti-GFP nanobody. (**ii**) Representative 2D classes of the BOS (fNOMO) complex. (**iii**) Data processing scheme in cryoSPARC v3.2-4.2.1 to identify a set of particles that correspond to the BOS (fNOMO) complex. After a round of blob picking and iterative rounds of 2D classification, 246,296 particles were used to generate 2D classes for selection. Then, 90,893 particles were used for Ab-initio reconstruction to generate the initial models. Using the model that best resembles the BOS (fNOMO) complex, rounds of heterogeneous refinement were performed on the previously mentioned 246,296 particles to identify the final set of 63,018 particles corresponding to our final map of BOS (fNOMO). **(C)** BOS (tNOMO). (**i**) Sample preparation of BOS (tNOMO). Left: Size exclusion chromatogram of BOS (tNOMO) sample after GFP-immunoprecipitation and TEV cleavage. UV absorbance at 280nm was recorded to track protein fractions that contained BOS (tNOMO) complex. Right: SDS-PAGE gel of the peak fraction containing purified BOS (tNOMO) complex, stained by Sypro Ruby. The components of the BOS (tNOMO) complex were present (TMEM147, NCLN, and tNOMO). Since the bait used for purification was TMEM147-GFP, a portion of endogenous full-length NOMO was also detected. (**ii**) Representative 2D classes of BOS (tNOMO) complex. (**iii**) Processing scheme used in cryoSPARC to identify particles that represent the BOS (tNOMO) complex. After an initial round of blob picking and iterative rounds of 2D classification, 319,567 particles were used to generate templates for template picking. Picked particles were put through iterative rounds of 2D classification and 1,265,788 particles were used to generate initial models to be used for multiple rounds of heterogeneous classification. 168,947 particles were subjected to 3D classification to identify the final set of 115,841 particles. The EM map corresponding to these particles was further refined using non-uniform refinement and sharpened with DeepEMhancer to yield the final map. (iv) Gold Standard Fourier Shell Correlation (GFSC) curves for the masked and unmasked half maps of the overall BOS (tNOMO) complex map. Nominal resolution of 3.6 Å was determined using gold standard FSC cutoff = 0.143. (v) Resolution estimation of the BOS (tNOMO) complex, generated from cryoSPARC.

**Figure S5.**
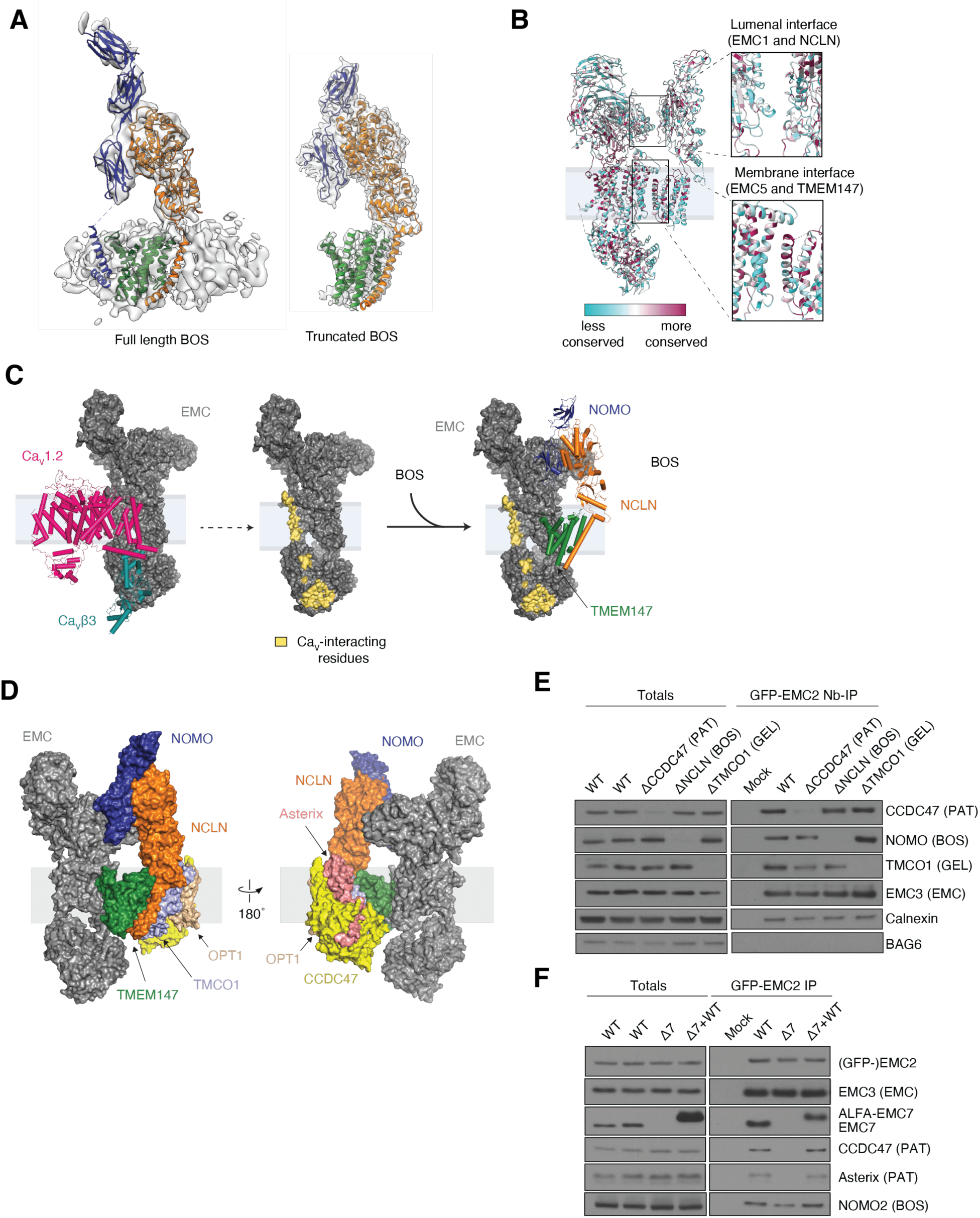
Analysis of the BOS complex and EMC • BOS (fNOMO) complex. (**A**) Comparison between the full-length BOS complex and truncated BOS complex model docked in their respective EM map. From the maps, we observed that the truncated BOS complex retained the overall shape of the full-length complex, although only the last 2 IgG-like domains are visible in the truncated BOS complex map. Here, NOMO is colored in blue, TMEM147 in green, and NCLN in orange. **(B)** Sequence conservation analysis of the EMC • BOS (fNOMO) complex. Left: Molecular model of EMC • BOS (fNOMO) with residues being colored by sequence conservation. Less conserved residues are in cyan and more conserved residues are in dark magenta. The sequence alignment contains sequences of EMC • BOS (fNOMO) from the following species: *Homo sapiens*, *Rattus norvegicus*, *Lygus hesperus*, *Callithrix jacchus*, *Chionoecetes opilio*, *Hydra vulgaris*, *Branchiostoma lanceolatum*, *Astyanax mexicanus*, *Castor canadensis*, *Pan troglodytes*, *Neovison vison*, *Macaca mulatta*, *Bactrocera latifrons*, *Bactrocera dorsalis*. Right: Inlets showing conservation of the lumenal interface (between EMC1 and NCLN) and membrane interface (between EMC5 and TMEM147). We observed a conserved patch near the lumenal membrane between EMC5 and TMEM147. **(C)** Comparison of the binding surfaces of EMC with either *Ca*V1.2/*Ca*Vβ3 complex or the BOS complex. The model was generated by aligning the EMC from its structure with *Ca*V1.2/*Ca*Vβ3 complex (PDB ID 8EOI) (Chen et al., 2023) or BOS complex (this study). Left: Cylinder cartoon representation of the *Ca*V1.2/*Ca*Vβ3 complex (pink and dark cyan)(Chen et al., 2023) and surface representation of the EMC (dark grey). Middle, the EMC residues that are interacting with *Ca*V1.2/*Ca*Vβ3 are highlighted in yellow. These residues were determined using the *InterfaceResidues.py* script in PyMOL. Right: Cylinder cartoon representation of the BOS complex (TMEM147 in green, NCLN in orange, and NOMO in blue) and surface representation of the EMC (dark grey) with the *Ca*V-interacting residues being highlighted to show that there is no overlap between the interacting interfaces with EMC by *Ca*V and the BOS complex. **(D)** Superposition of EMC • BOS (fNOMO) complex with the other components of the ‘ER translocon’ at Sec61. The model was generated by aligning TMEM147 from EMC • fNOMO (this study) and the ER translocon (PDB ID 6W6L) (McGilvray et al., 2020). Sec61 and CCDC47’s latch helices were removed from this model for clarity. EMC occupies the same position as Sec61 at the translocon. **(E)** HEK 293T cells (WT, ΔCCDC47, ΔNCLN, or ΔTMCO1) stably expressing GFP-EMC2 were subjected to anti-GFP nanobody purification under native conditions. Samples were analyzed by SDS-PAGE and western blotting. Note that the interaction of the EMC with individual multipass translocon components is independent of the EMC’s interactions other multipass translocon components. **(F)** EMC7 is indispensable for interaction of the EMC with multipass translocon components. WT or EMC7 KO cells (Δ7) stably expressing GFP-EMC2 were transduced with WT EMC7 or a BFP control and subjected to anti-GFP nanobody purification under native conditions. Samples were analyzed by SDS-PAGE and western blotting.

**Figure S6.**
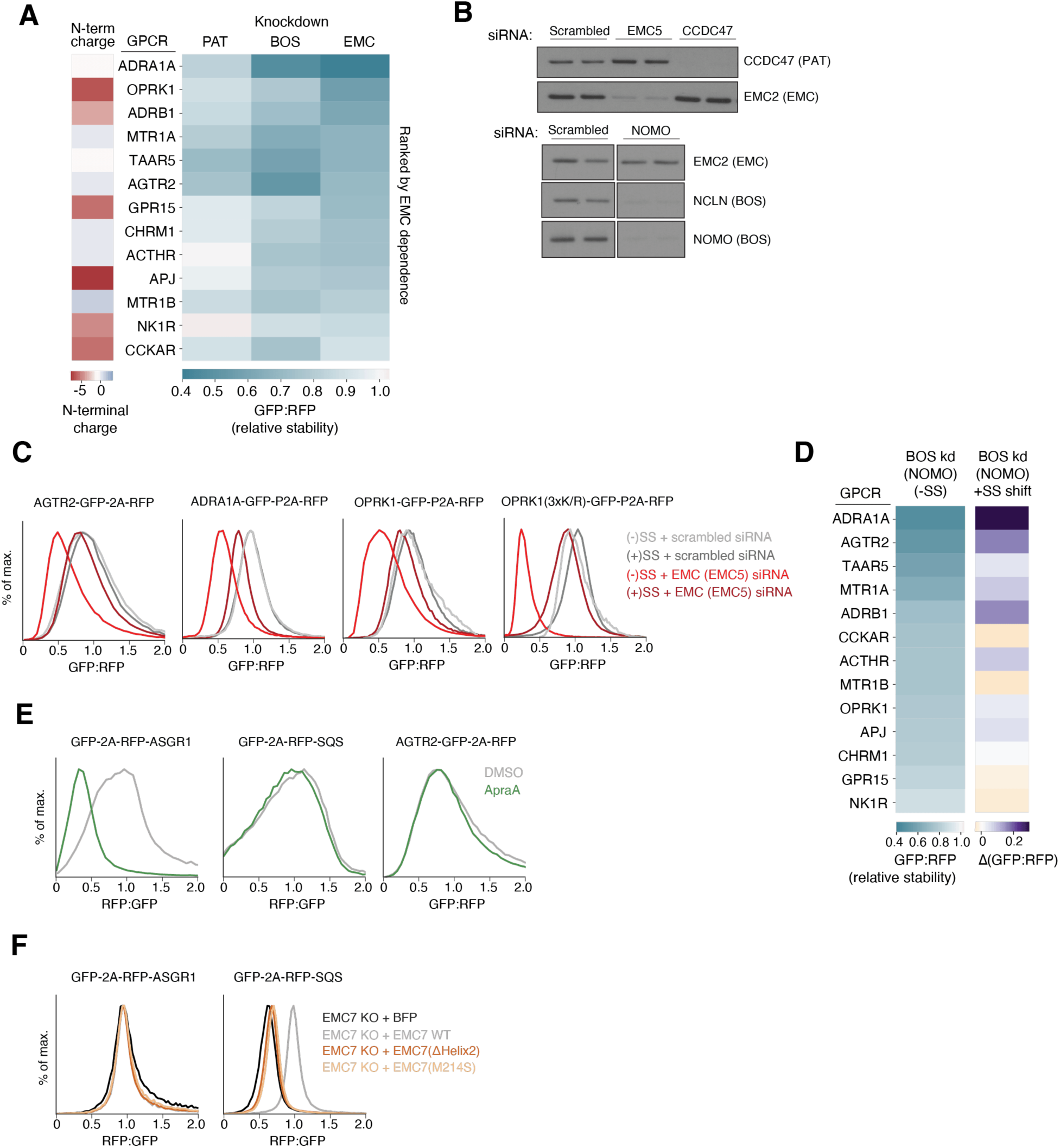
Differential effects of biogenesis factors on diverse GPCR substrates. **(A)** A panel of GPCRs was assayed for EMC, BOS, or PAT dependence, as described in Figure 6. (Right) A heatmap displays the relative stability of each GPCR, ranked by EMC dependence. (Left) The net charge of the N-terminal soluble domain of each GPCR is indicated by the heatmap. **(B)** Validation of knockdown experiments in RPE1 cells using siRNAs targeting EMC5, CCDC47, NOMO used in Figures 4 and 6. Cells were harvested and analyzed by SDS-PAGE and western blotting. **(C)** Flow cytometry assay as in Figure 4C and 6C, with EMC5 knockdown. **(D)** Bypassing the EMC for insertion of TMD1 of GPCRs reduces the dependence on the BOS complex. GPCR reporter constructs were generated using the GFP-2A-RFP cassette, with and without a signal sequence (SS) and T4 lysozyme fusion to the N-terminus of each GPCR. RPE1 cells were transduced with these reporters after siRNA knockdown of NOMO or a non-targeting control. (Left) A heatmap shows the relative stability of GPCRs without a signal sequence and is ranked by BOS dependence. (Right) A heatmap shows the change in stability between GPCRs containing the appended signal sequence and WT GPCRs. Darker purple indicates a positive shift in stability of the signal sequence-containing GPCR compared to the WT GPCR. **(E)** As in Figure 6E, for ASGR1, SQS, and AGTR2. **(F)** As in Figure 6F, for ASGR1 and SQS.

**Figure S7.**
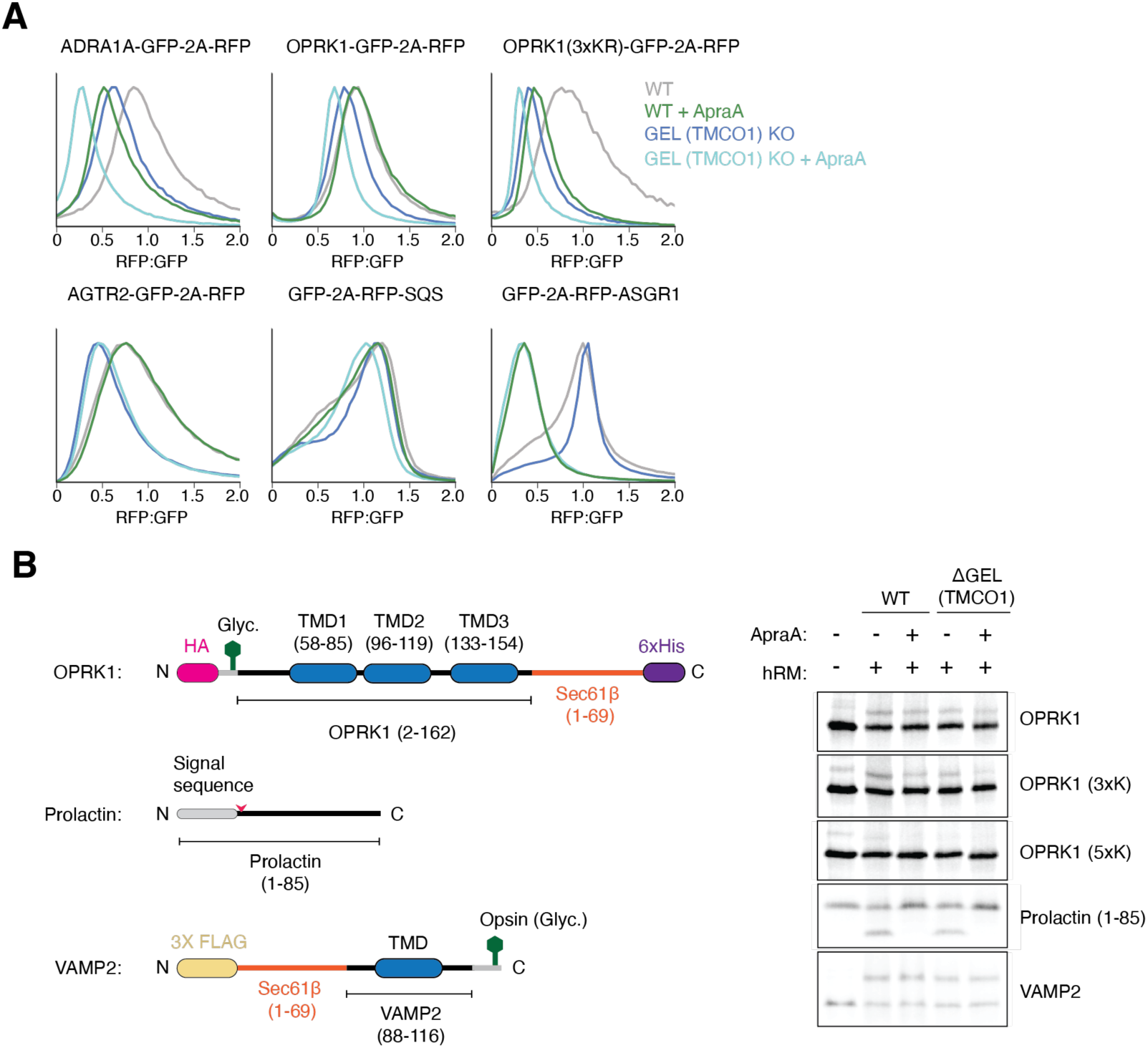
Substrates are routed through TMCO1 and Sec61 when unable to access EMC for insertion. **(A)** Histograms as in Figure 6D,E, for WT and TMCO1 KO cells treated with Apratoxin A or a DMSO control. **(B)** *In vitro* translation of OPRK1 (wildtype, 3xK, or 5xK), Prolactin, or VAMP2. (Left) Schematic of constructs used in the translation reactions. The OPRK1 3xK and 5xK variants contain additional positive charges in the N-terminal soluble domain, as described in Figure 6B. (Right) Translations were performed in RRL in the presence of either 1 μM Apratoxin A in 0.5% DMSO or a 0.5% DMSO control and in hRMs derived from WT or TMCO1 KO cells. Insertion was determined by the fraction of glycosylated substrate (OPRK1 variants and VAMP2) or by signal sequence cleavage (Prolactin). In vitro insertion reactions tend to result in smaller phenotypes than those observed in cells, in part due to the absence of competing off-pathway reactions, including protein quality control and surveillance. Additionally, the time scales of an in vitro insertion reaction are beyond the physiologic time scale in cells, which gives substrates near-infinite time to be inserted into the ER microsomes.

**Table S5.**
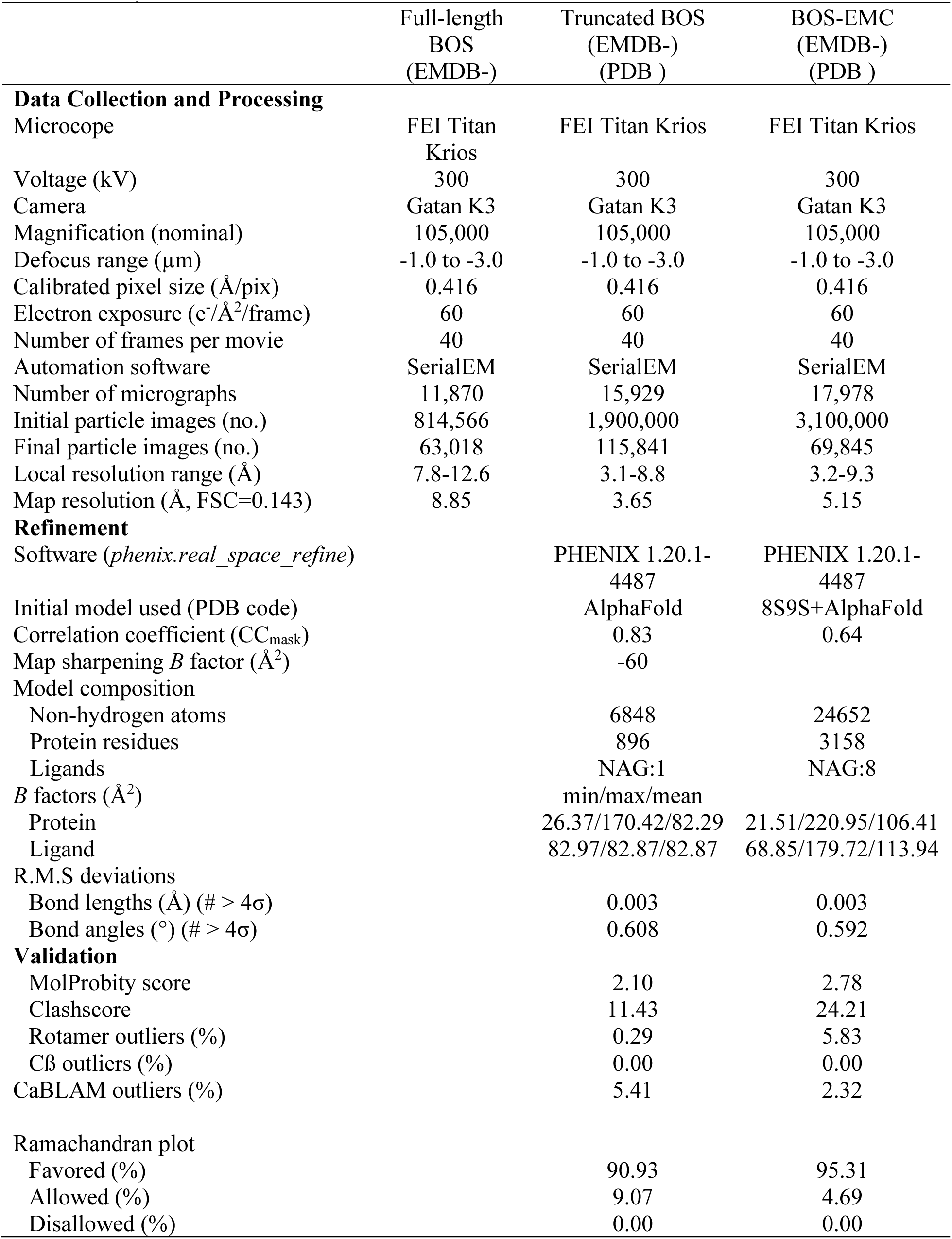
CryoEM Data Collection, Refinement, and Validation Statistics.

## Supplementary File Legends

Supplemental Table 1. Genome-wide FACS screens with non-targeting and EMC2 dual libraries for AGTR2

Supplemental Table 2. Genome-wide FACS screens with single guide libraries for SARS-CoV-2 M

Supplemental Table 3. Genome-wide FACS screens with single guide libraries for SARS-CoV-2 ORF3a

Supplemental Table 4. Mass spectrometry of the EMC

